# Choanoflagellates discriminate among *Pseudomonas aeruginosa* alkylquinolones during pathogen response

**DOI:** 10.64898/2026.11.26.768673

**Authors:** Jan Pauline Gabriel, Collin McNairy, Marie Dayras, Chia-Chi Peng, Vicki Deng, Francois-Xavier Meyer, Christine Beemelmanns, Arielle Woznica

**Author notes:** J.P.G., C.M., and M.D. contributed equally to this work. In memory of Jon Clardy.

## Abstract

Choanoflagellates are emerging models for studying host-pathogen interactions, and as the closest living relatives of animals, offer a window into the evolution of immune defense against bacteria. However, investigating mechanisms of pathogen recognition and response in choanoflagellates remains challenging in the absence of defined bacterial effectors. By studying the interaction between the pathogen *Pseudomonas aeruginosa* and the choanoflagellate *Monosiga brevicollis*, we identify alkylquinolone HNQ (2-nonyl-4-quinolone) as a potent effector that reduces host viability. Structure–activity studies reveal that both the nine-carbon side chain and the unmodified 4-quinolone core contribute to HNQ’s potency. HNQ is enriched in outer membrane vesicles, suggesting a route for its delivery in aquatic environments. In addition, HNQ reduces the viability of diverse choanoflagellate species and human cells, indicating that its effector activity may extend across evolutionarily distant hosts. Notably, *M. brevicollis* distinguishes HNQ, a molecule produced primarily by *P. aeruginosa*, from closely related hydroxylated compounds and mounts a selective response involving MbABCG1, a novel ABCG transporter that modulates sensitivity to HNQ. Together, these findings identify HNQ as a broadly active *P. aeruginosa* effector and show that choanoflagellates distinguish among closely related metabolites produced by a pathogen, providing a foundation for understanding the molecular basis of pathogen recognition and response.

**SIGNIFICANCE:** *Pseudomonas aeruginosa*, a pathogen of diverse eukaryotes from amoebae to humans, secretes dozens of related molecules called alkylquinolones (AQs). By studying pathogen response in choanoflagellates, the closest living relatives of animals, we identify HNQ, an abundant but understudied AQ, as an effector that reduces the viability of choanoflagellates and human cells. We find that choanoflagellates have distinct responses to structurally similar alkylquinolones, and identify a cell-surface transporter that promotes survival upon HNQ exposure. Because HNQ is made predominately by *P. aeruginosa*, our findings illustrate how choanoflagellates can recognize a specific bacterial pathogen among diverse bacteria, provide a new probe for dissecting pathogen response in choanoflagellates, and identify an effector that may prove broadly relevant to *P. aeruginosa* pathogenesis.

## INTRODUCTION

Choanoflagellates are an emerging system for investigating host-pathogen interactions. As the closest relatives of animals (Figure 1A), choanoflagellates uniquely share many cell biological and genetic features with animals (1, 2), and studying choanoflagellates has already yielded insights into the evolution of animal immune signaling (2, 3). The ecology of choanoflagellates provides a further compelling reason to study their responses to pathogens. These unicellular or facultatively multicellular bacterivores live in bacteria-dominated aquatic environments, where they encounter bacteria as both prey and potential pathogens (4). Whereas many animal cells are shielded from direct contact with bacteria, choanoflagellates continually encounter bacteria and their secreted products at the cell surface, and they likely rely on specific molecular cues to distinguish among bacteria (5–8). Nonetheless, the bacterial molecules that elicit immune responses in choanoflagellates remain almost entirely unknown. Identifying such effectors would provide critical molecular tools for dissecting how choanoflagellates recognize and respond to pathogens.

**Figure 1.**
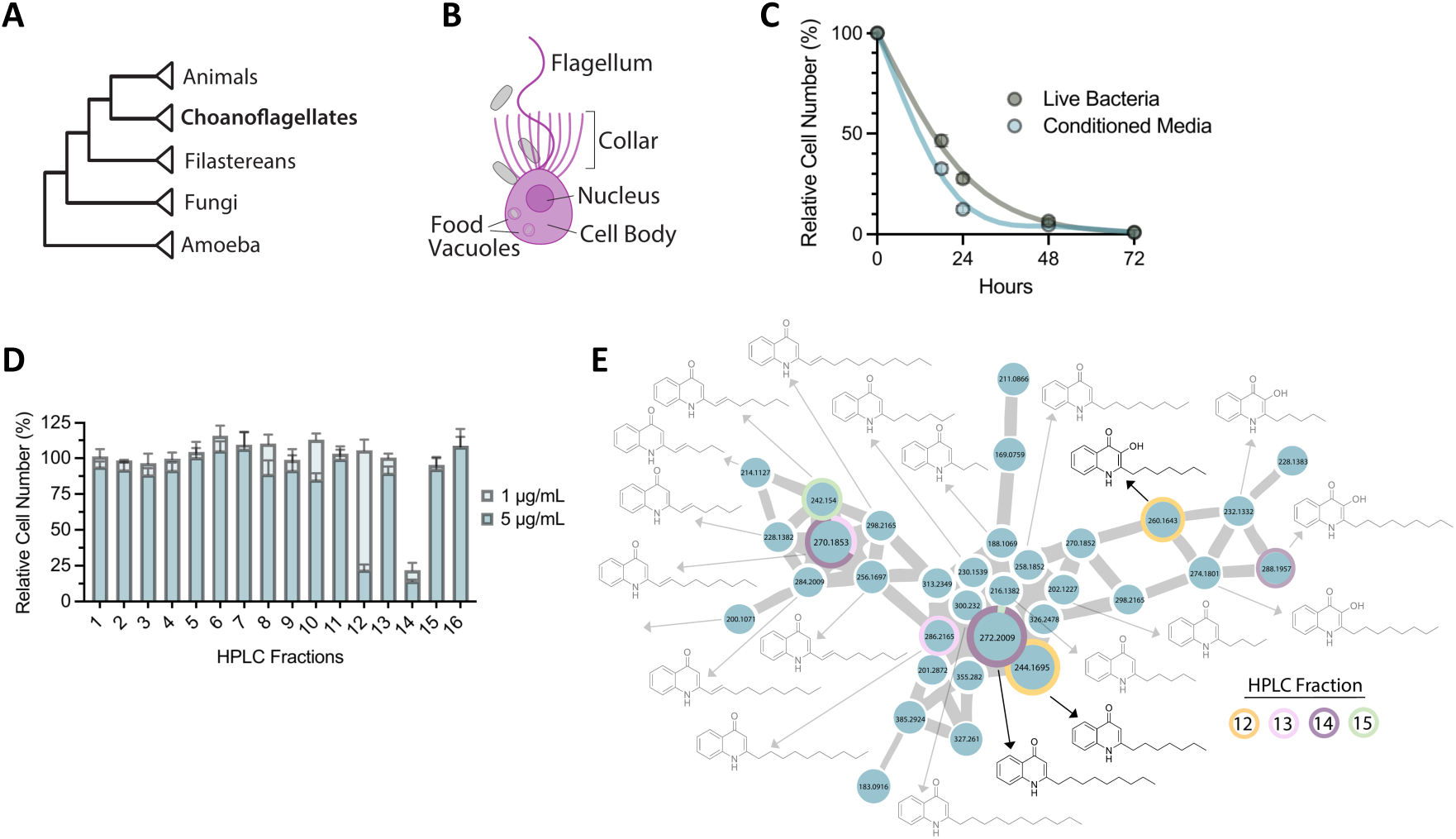
Activity-guided fractionation reveals that secreted *P. aeruginosa* alkylquinolones reduce *M. brevicollis* viability **A.** Phylogeny showing that choanoflagellates are the sister group of animals. Their close relationship to animals makes them a simplified yet powerful model for studying host-pathogen interactions. **B.** Schematic of *M. brevicollis*, a motile unicellular choanoflagellate. *M. brevicollis* bears a single apical flagellum that is surrounded by a feeding collar composed of actin-filled microvilli. Prey bacteria are captured at the feeding collar and internalized into food vacuoles by phagocytosis. **C.** Secreted *Pseudomonas aeruginosa* (PAO1) factors are sufficient to reduce *M. brevicollis* viability. *M. brevicollis* cell density was quantified at the indicated time points following exposure to PAO1 live bacteria (MOI 50) or PAO1 cell-free conditioned media (5% vol/vol). Relative cell number (%) was calculated by comparing samples treated with PAO1 live bacteria or conditioned media to samples treated with the feeding bacterium *P. shioyasakiensis* (live bacteria or conditioned media, respectively). PAO1 live bacteria and conditioned media similarly reduced *M. brevicollis* viability, demonstrating that secreted factors alone are sufficient. Data represent mean +/- SD for three biological replicates. **D.** Activity-guided fractionation of PAO1 conditioned media identifies bioactive compounds for further study. PAO1 (*rhlA^-^*) conditioned media organic extract was fractionated using reversed-phase High Performance Liquid Chromatography (HPLC). The resulting 16 fractions were dried, resuspended in DMSO, and added to *M. brevicollis* at a final concentration of 1 or 5 µg/mL. *M. brevicollis* cell density was quantified after 22 hours, and relative cell number (%) was calculated for each sample against a DMSO control. While Fractions 12 and 14 reduced viability, Fraction 14 was the most potent, and the only fraction to reduce viability at the lower concentrations. Data represent mean +/- SD for three biological replicates. **E.** Bioactive Fractions 12 and 14 contain chemically distinct alkylquinolone congeners. Expanded view of a GNPS feature-based molecular network cluster of quinolones from the *P. aeruginosa* PAO1 (*rhlA^-^*) organic extract. HNQ (2-nonyl-4-quinolone) is the predominant congener in Fraction 14, whereas HHQ (2-heptyl-4-quinolone) and PQS (2-heptyl-3-hydroxy-4-quinolone) are the predominant congeners in Fraction 12. Annotated compounds in black (PQS, HHQ, and HNQ) were confirmed by comparison to synthetic standards or by structural elucidation after isolation (Level 1 confidence). Compounds in grey were annotated with a confidence level of 2 or 3 (MS/MS spectral matching against spectral libraries or comparison with spectra from the same compound family). Major ions (intensity > 10*^8^*) are displayed as pie charts, with colored slices representing their relative contribution across Fractions 12-15. Node size is proportional to feature abundance (peak area), and edge width reflects MS/MS spectral similarity (cosine score).

We previously established *Monosiga brevicollis* as an experimentally tractable model for studying choanoflagellate immune response, and found that it is susceptible to *Pseudomonas aeruginosa* (2). *P. aeruginosa* is a ubiquitous environmental bacterium that exerts pathogenic effects on a broad range of eukaryotic hosts, from amoebae to mammals (9). This versatility derives from its large arsenal of secreted proteins and secondary metabolites, including the 2-alkyl-4(1H)-quinolones (AQs) (9). The best-studied *P. aeruginosa* quinolones are the quorum-sensing molecule PQS (*Pseudomonas* Quinolone Signal) and its biosynthetic precursor HHQ (2-heptyl-4(1H)-quinolone), which regulate interbacterial communication, biofilm formation, and the production of virulence factors (10–17). PQS and HHQ have also been shown to directly modulate eukaryotic cell signaling and viability (14, 18–24). However, *P. aeruginosa* produces over fifty other quinolones (11, 17, 25) whose functions during interkingdom and host-pathogen interactions remain largely unknown.

We recently showed that secreted *P. aeruginosa* factors elicit an immediate and robust transcriptional response in *M. brevicollis* and that prolonged exposure results in cell death (Figure 1C). However, despite screening many candidate strains carrying mutations in known virulence or secretory genes, we were unable to resolve the *P. aeruginosa* effectors responsible for reduced viability. Here, we use activity-guided fraction to identify HNQ (2-nonyl-4(1H)-quinolone), an abundant but understudied *P. aeruginosa* alkylquinolone, as a potent effector that reduces *M. brevicollis* viability. Through structure–activity analyses, we define the chemical features required that contribute to HNQ activity, and show that non-hydroxylated AQs are necessary for *P. aeruginosa* to exert its full effect on *M. brevicollis* viability. HNQ also reduces the viability of diverse choanoflagellate species and human cells, and is enriched in bacterial outer membrane vesicles, providing a plausible route for delivering this lipophilic effector in both aquatic environments and infection contexts. Finally, we identify a choanoflagellate ABCG transporter, MbABCG1, whose abundance increases specifically in response to HNQ and related non-hydroxylated AQs and whose expression modulates cellular sensitivity to HNQ. Together, our findings establish HNQ as a previously overlooked *P. aeruginosa* effector relevant during host-pathogen interactions and provide a molecular foothold for understanding how choanoflagellates recognize and respond to pathogens amid a complex bacterial environment.

## RESULTS

### Secreted *P. aeruginosa* alkylquinolones reduce *M. brevicollis* viability

After determining that the activity of *P. aeruginosa* conditioned media is fully retained after organic extraction, we fractionated the extract using reversed-phase High Performance Liquid-Chromatography (HPLC) (Figure S1A). Screening the resulting HPLC fractions across several concentrations identified two fractions, 12 and 14, that reduced *M. brevicollis* survival (Figure 1D). Fraction 14 was the most potent, and the only fraction that reduced cell viability at low concentrations (≤1 µg/mL). Analyzing all HPLC fractions by UHPLC-HRMS (Ultra-High Performance Liquid Chromatography-High Resolution Mass Spectrometry), and performing dereplication with metabolomic databases(26, 27), identified rhamnolipids and quinolones as the major components of the active fractions. At this stage, because we had previously determined that rhamnolipid biosynthesis is not required for *P. aeruginosa* cytotoxicity towards *M. brevicollis*(2), we could exclude rhamnolipids as key effectors. Therefore, after confirming that the secreted metabolomic profiles of parental (mPAO1) and rhamnolipid-deficient (rhlA::Tn) strains are otherwise similar under our growth conditions (Figure S1A), we chose to streamline subsequent purifications by using *P. aeruginosa* rhlA::Tn conditioned media (Figure 1D).

We performed further bioactivity-guided fractionation of enriched *P. aeruginosa* rhlA::Tn extracts using a combination of HPLC, UHPLC-HRMS, and dereplication workflows. This again identified quinolone features as the most prominent in the active fractions, which we confirmed by comparing the molecular ion features of our putative identifications to commercial synthetic standards (Figure 1E). Indeed, comparative LC-MS/MS analysis revealed that the bioactive fractions 12 and 14, and the surrounding but inactive fractions, contained 2-alkyl-4-quinolones (AQs), a class of secondary metabolite synthesized primarily by *Pseudomonas* bacteria (Figure 1E)(17, 25).

Although the bioactive fractions contained structurally related scaffolds, each fraction was composed of chemically distinct AQ congeners (Figure 1E; Figure S1B). The more polar fraction (Fraction 12) contained the quorum-sensing molecules HHQ and PQS, both bearing seven-carbon side chains based on their MS/MS fragmentation pattern and comparison with standards (10, 11, 13–15). The less polar fraction (Fraction 14) contained three structurally related AQs with nine-carbon side chains, including 2-nonyl-4-quinolone (HNQ), an abundant *P. aeruginosa* AQ that has remained largely unstudied during host-pathogen interactions (Figure 1E)(28, 29).

### Structure-activity analysis reveals HNQ as a potent effector

To corroborate our metabolomics data and directly assess the contribution of each compound to the observed activity of fractions 12 and 14, we used commercial and synthesized standards(30–32) of candidate AQs to assess whether they are sufficient to reduce *M. brevicollis* viability (Figure 2). We tested each compound at a range of concentrations, and determined that HNQ, the predominant component of fraction 14, is the most potent. Structure-activity analysis of HNQ and congeners further revealed that AQ bioactivity is dependent on both alkyl chain length and the 4-quinolone core (Figure 2A-D). Shortening or lengthening the alkyl chain of HNQ (C_9_) reduced potency: for example, HHQ (C_7_) differs from HNQ by only two carbons yet is 6-fold less potent, and analogs with chains longer than nine carbons exhibit drastically diminished activity (Figure 2B). Hydroxylating HNQ at the 3’ position (to yield 2-nonyl-3-hydroxy-4-quinolone) decreased both potency and efficacy (Figure 2C). Modifying the 4-quinolone scaffold of HHQ similarly reduced bioactivity: PQS has diminished maximal activity compared to HHQ, and HQNO, which reportedly disrupts electron transport in some eukaryotes(33), has little effect on *M. brevicollis* viability (Figure. 2D). Together, these experiments establish HNQ as a *P. aeruginosa* effector capable of modulating eukaryotic cell viability, define structural features critical for the bioactivity of HNQ towards *M. brevicollis*, and provide insight into the specificity of HNQ as an agonist.

**Figure 2.**
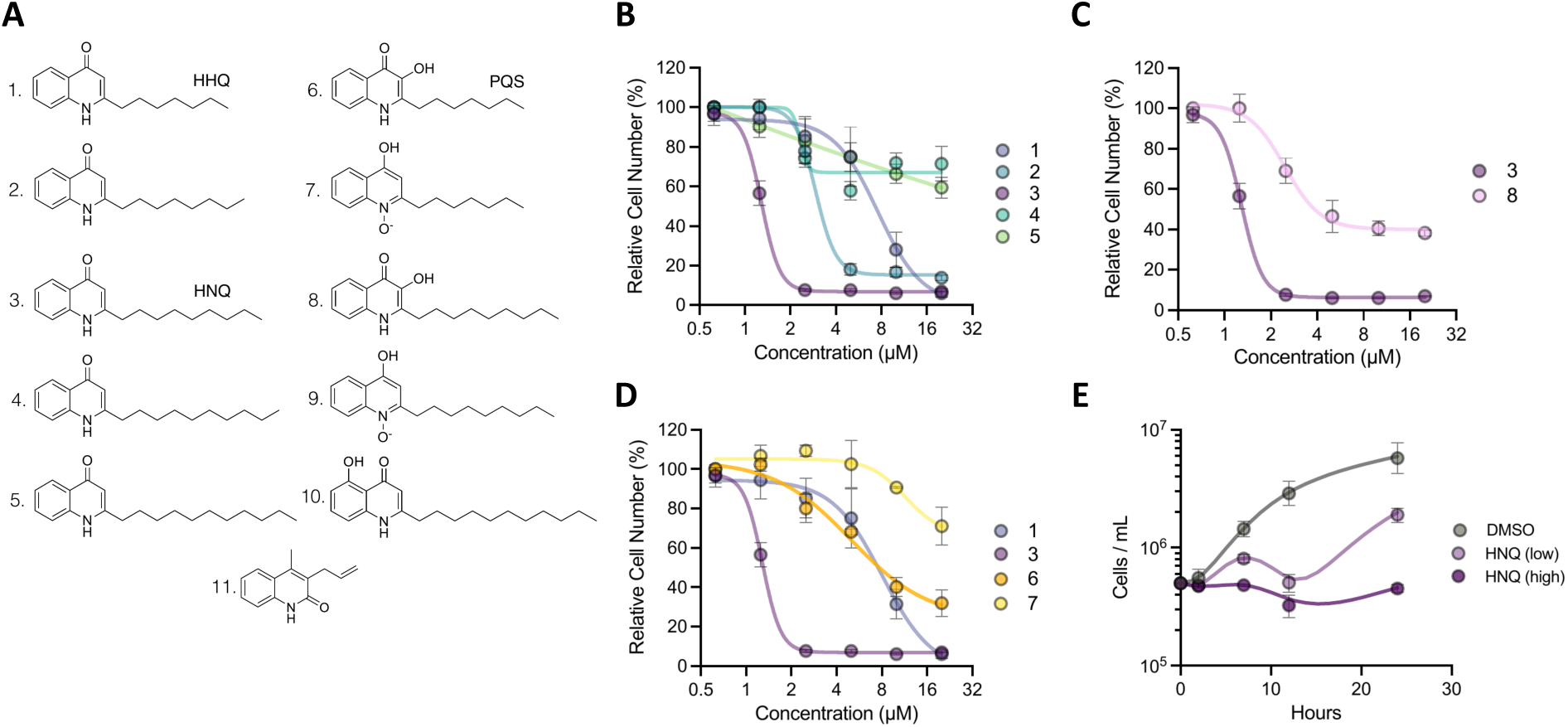
HNQ is a potent effector whose bioactivity depends on alkyl chain length and the 4-quinolone core **A.** Structures of the synthetic alkylquinolones (compounds 1-11) used in the structure-activity analyses, including HHQ (1), HNQ (3), and PQS (6). Compounds vary systematically in alkyl side-chain length and in modifications to the 4-quinolone core (e.g., 3’-hydroxylation, N-oxidation), enabling the contributions of these features to be tested independently. For all dose-response experiments in B-D, *M. brevicollis* was treated with the indicated compounds across a range of concentrations, and viability was evaluated after 22 hours by quantifying relative cell number (%) against a DMSO control. Data represent mean +/- SD for four biological replicates. **B.** HNQ is more potent than congeners with a shorter or longer alkyl chain. Dose-response curves comparing HNQ (3) with alkylquinolones bearing shorter (compounds 1 & 2) or longer (compounds 4 & 5) side chains. **C.** Modifying the 4-quinolone core of HNQ decreases its potency and efficacy. Dose-response curves comparing HNQ (compound 3) with its 3’-hydroxylated derivative, 2-nonyl-3-hydroxy-4-quinolone (compound 8). **D.** Bioactivity of HNQ compared to the well-characterized alkylquinolones HHQ (compound 1), PQS (compound 6), and HQNO (compound 7). **E.** Time course of HNQ exposure. *M. brevicollis* was treated with a half-maximal (“low”; 1.25 uM) or maximal (“high”; 2.5 uM) inhibitory concentration of HNQ, or with a DMSO control, and cell density (cells/mL) was quantified at the indicated time points over 24 hours. At the low dose, cells continued to proliferate for roughly one cell cycle (∼5-6 hours) before cell numbers declined and then partially recovered. At the high dose, initial proliferation was suppressed but cell death was not observed at early time points. Data represent mean +/- SD for three biological replicates.

During infection, eukaryotic hosts encounter multiple alkylquinolones simultaneously. Moreover, concentrations of quorum sensing molecules HHQ and PQS, and likely other alkylquinolones, fluctuate throughout *P. aeruginosa* growth and in response to changing environmental conditions(11, 34–36). Therefore, we examined how co-exposure with HNQ affects the potency of HHQ and PQS. We treated *M. brevicollis* with a sublethal concentration of HNQ (0.625 µM) and titrated the concentration of HHQ or PQS (Figure S2A-C). Co-treatment with HNQ additively enhanced the potency of HHQ (Figure S2A,B). In contrast, co-treatment with HNQ did not affect the potency or efficacy of PQS (Figure S2C). These results raise the possibility that non-hydroxylated (HHQ, HNQ) and hydroxylated (PQS) alkylquinolones are perceived through distinct M. brevicollis receptors.

HNQ has not been previously described to reduce eukaryotic cell viability. Therefore, to investigate the kinetics of HNQ-induced responses in *M. brevicollis*, we monitored cell behavior and quantified growth at multiple time points following treatment with either an intermediate (1.25 µM, which is slightly below the calculated LD50) or high (2.5 µM) concentration of HNQ. For at least 5 hours post-treatment, flagellar beating remained largely unaffected, and cells continued to feed and swim (Movies S1, S2). Consistent with these observations, growth curves showed that cell populations treated with 1.25 µM HNQ continued to proliferate for about one cell cycle – roughly 5-6 hours for *M. brevicollis* – before cell numbers began to decline (Figure 2E). At the higher HNQ dose, initial cell proliferation was reduced but, notably, cell death did not occur at earlier time points (Figure 2E). Growth curves further revealed that, following a period of cell death, populations exposed to lower HNQ concentrations largely recovered as surviving cells resumed proliferation. Importantly, choanoflagellate cell death was not caused by starvation, as treating *M. brevicollis* with high concentrations of HNQ did not kill co-cultured feeding bacteria (Table 1). These kinetic data, together with our structure-function analysis, lead us to hypothesize that HNQ modulates *M. brevicollis* signaling through interactions with a specific host factor, and that one downstream outcome of HNQ-induced signaling is regulated cell death.

### Non-hydroxylated alkylquinolones are necessary for the cytotoxic activity of *P. aeruginosa* conditioned media

Because *P. aeruginosa* secretes several factors that are cytotoxic towards other eukaryotes(37, 38), we next investigated whether AQs are necessary for conditioned media to exert its full effect on *M. brevicollis* viability. We quantified choanoflagellate survival following treatment with conditioned media isolated from *P. aeruginosa* strains carrying mutations in quinolone biosynthesis genes (Figure 3A,B)(39–41). Because PQS itself regulates the production and secretion of virulence factors at the onset of stationary phase(14, 16, 42), conditioned media from mid-exponential phase cultures was harvested to avoid confounding effects.(37)

**Figure 3.**
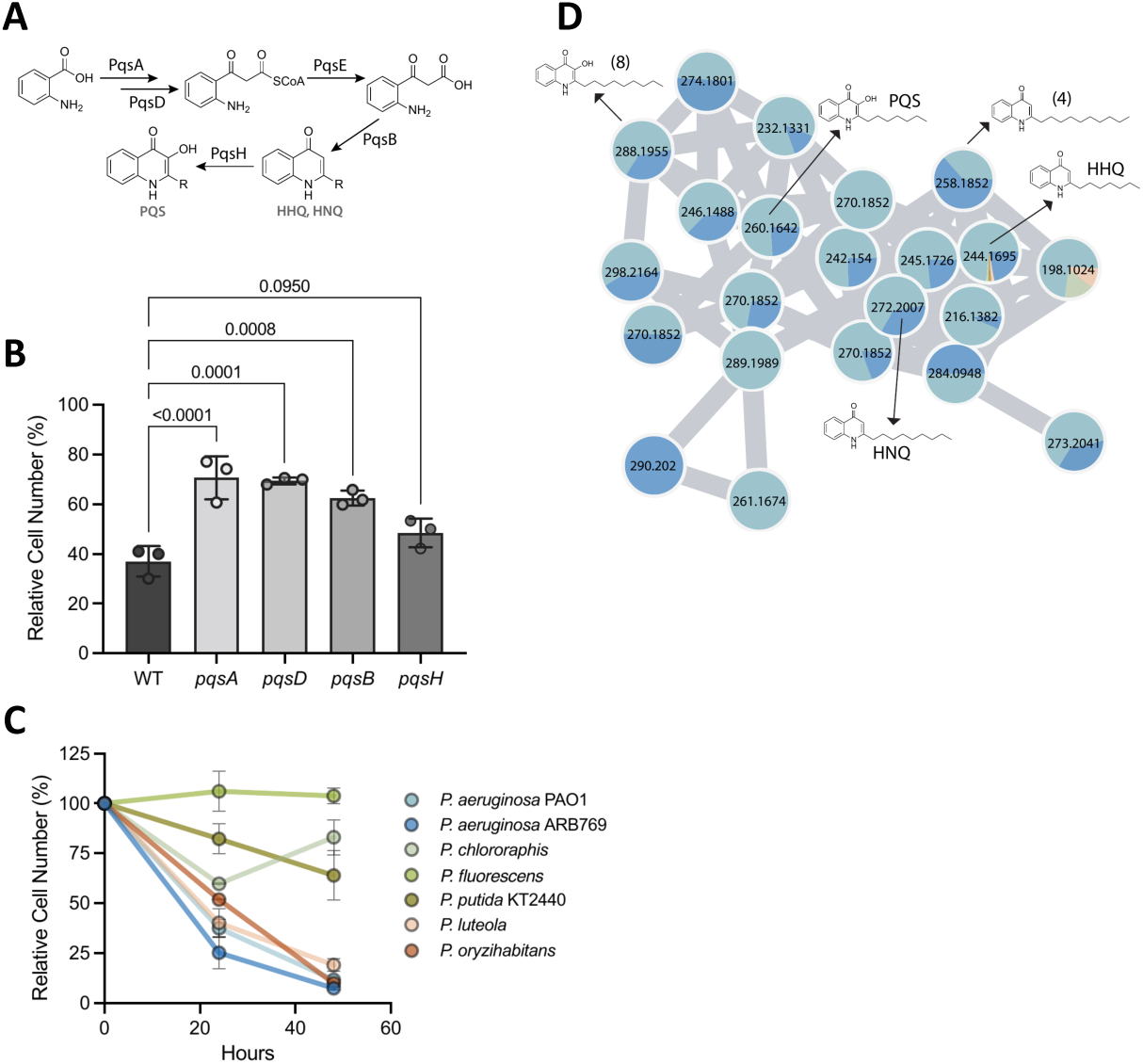
Non-hydroxylated alkylquinolones contribute to the cytotoxicity of *P. aeruginosa*. **A.** Simplified schematic of the *P. aeruginosa* alkylquinolone biosynthesis pathway. Anthranilate is activated and condensed by PqsA and PqsD, and the product is processed via PqsE and PqsBC to yield the non-hydroxylated 2-alkyl-4-quinolones HHQ and HNQ. PqsH then 3-hydroxylates these congeners to produce PQS (and its C9 analog). This organization means that *pqsA*, *pqsD*, and *pqsB* mutants fail to make any AQs, whereas a *pqsH* mutant still makes HHQ and HNQ but cannot 3-hydroxylate them. R, alkyl side chain. **B.** Non-hydroxylated AQ biosynthesis is required for the cytotoxic activity of *P. aeruginosa* conditioned media. *M. brevicollis* was treated with conditioned media (5% vol/vol) harvested from mid-exponential phase cultures (OD600 = 1.2) of the parental strain (WT; mPAO1) or of *pqsA*, *pqsD*, *pqsB*, or *pqsH* transposon mutants, and relative cell number (%) was quantified after 22 hours. Relative cell number was calculated by comparison to treatment with *P. shioyasakiensis* conditioned media. Each point represents one biological replicate; bars represent mean +/- SD. Statistical analysis was performed using a RM one-way ANOVA. Shown are the adjusted P-values from a Dunnett’s multiple comparison test. **C.** *Pseudomonas* species vary in their cytotoxicity toward *M. brevicollis*. Relative cell number (%) of *M. brevicollis* over 48 hours following exposure to conditioned media isolated from the indicated *Pseudomonas* species. Both *P. aeruginosa* strains (PAO1, ARB769) strongly reduced viability; among the non-*aeruginosa* species, *P. chlororaphis*, *P. fluorescens*, and *P. putida* had little effect, whereas *P. luteola* and *P. oryzihabitans* reduced viability by 48 hours despite not producing detectable AQs. Data represent mean +/- SD for three biological replicates. **D.** *P. aeruginosa* uniquely synthesizes a chemically diverse array of alkylquinolones. Feature-based molecular network of secreted metabolites from *P. aeruginosa* PAO1, clinical *P. aeruginosa* isolate ARB769, and five diverse *Pseudomonas* species. Nodes represent MS features (annotated with precursor *m/z*); pie charts indicate the relative contribution of each species to a given feature. AQs tested in structure-function analysis (HNQ, PQS, HHQ, and congeners) are indicated. HNQ is detected exclusively in *P. aeruginosa*. HHQ is present at very low abundance in the other *Pseudomonas* species.

*M. brevicollis* was significantly less susceptible to conditioned media from *pqsA, pqsD,* and *pqsB* mutant strains, which we confirmed produce no detectable AQs, than to conditioned media from the parental mPAO1 strain (Figure 3B; Table 2; Table 3). In contrast, the effect of conditioned media from the *pqsH* mutant did not differ significantly from that of the parental strain (Figure 3B). This is consistent with the role of PqsH, which is responsible solely for the 3-hydroxylation of AQs(11). Moreover, the *pqsH* mutant still produced HHQ, HNQ, and a small amount of PQS (Table 3). These data suggest that non-hydroxylated AQs, including HNQ, substantially contribute to the pathogenic effects of *P. aeruginosa* conditioned media.

We observed that AQ-deficient *P. aeruginosa* still slightly restricted *M. brevicollis* growth compared to non-pathogenic control bacteria (Figure S2D), suggesting that additional secreted factors may contribute to cytotoxicity. Diketopiperazines, a class of cyclic dipeptides with reported cytotoxic and antiproliferative effects on human cells(43, 44), were enriched in inactive HPLC fractions. However, these molecules were not bioactive alone or in combination with HNQ (Figure S2E, F), and the identity of these additional factors remains to be determined.

### *P. aeruginosa* has an expanded AQ repertoire

We next asked whether other *Pseudomonas* species exert similar pathogenic effects on *M. brevicollis.* Metabolomic studies of *Pseudomonas* alkylquinolones have primarily focused on *P. aeruginosa*, and the extent to which other *Pseudomonas* species synthesize AQs is unclear. Therefore, we characterized the secreted metabolite profiles of a clinical *P. aeruginosa* strain alongside those of five diverse *Pseudomonas* species.

Feature-based molecular networking revealed that only *P. aeruginosa* synthesized a chemically diverse array of AQ structures, with both *P. aeruginosa* strains PAO1 and clinical isolate ARB769 producing a broad spectrum of quinolone derivatives (Figure 3D). Notably, HNQ was exclusively detected in the *P. aeruginosa* strains, suggesting that it is not a broadly produced *Pseudomonas* metabolite. As expected from its metabolic profile, *P. aeruginosa* strain ARB769 reduced *M. brevicollis* viability similarly to PAO1 (Figure 3C).

We next asked whether other *Pseudomonas* species, which produce a limited diversity and abundance of AQs, are cytotoxic towards choanoflagellates. The five non-*aeruginosa* species variably impacted choanoflagellate viability: three species (*P. chlororaphis*, *P. fluorescens*, *P. putida*) had little effect on choanoflagellate growth, whereas *P. luteola* and *P. oryzihabitans* reduced *M. brevicollis* viability after 48 hours (Figure 3C). We assessed global metabolomic differences by principal component analysis (PCA), which revealed clear species-level separation (Figure S3F). Hierarchical clustering likewise confirmed strong intra-species reproducibility and indicated that only *P. luteola* and *P. oryzihabitans* have partially overlapping metabolomes (Figure S3G); however, because neither species produces AQs, their cytotoxicity is AQ-independent. Together, these data indicate that the expanded AQ repertoire of *P. aeruginosa* differentiates its metabolomic profile from other *Pseudomonas* species and may enhance its pathogenicity towards a broader range of eukaryotes, including choanoflagellates.

### Diverse choanoflagellate species and human cells are susceptible to HNQ

Given that *P. aeruginosa* thrives across a range of ecological niches and exerts pathogenic effects on diverse eukaryotes(9), we investigated whether HNQ functions as a virulence factor in other choanoflagellate and human host contexts. Choanoflagellates are genetically diverse and inhabit nearly all aquatic environments (1, 4). Therefore, to determine if HNQ affects the viability of choanoflagellate species beyond *M. brevicollis*, we selected three additional species for testing, each thriving under distinct environmental conditions and co-cultured with different bacterial communities (Figure 4A, 4B)(45–48). All three species were susceptible to HNQ: *C. flexa* showed comparable sensitivity to *M. brevicollis*, while *S. rosetta* and *M. fluctuans* were approximately twice as sensitive. To investigate if the structural requirements for AQ activity are conserved across choanoflagellate species, we also compared the potency of HNQ and HHQ for each species. HNQ was 5-7 fold more potent than HHQ across all species tested (Figure S3A). The potency and specificity of HNQ-induced responses in diverse choanoflagellates raises the possibility that HNQ targets a conserved choanoflagellate receptor.

**Figure 4.**
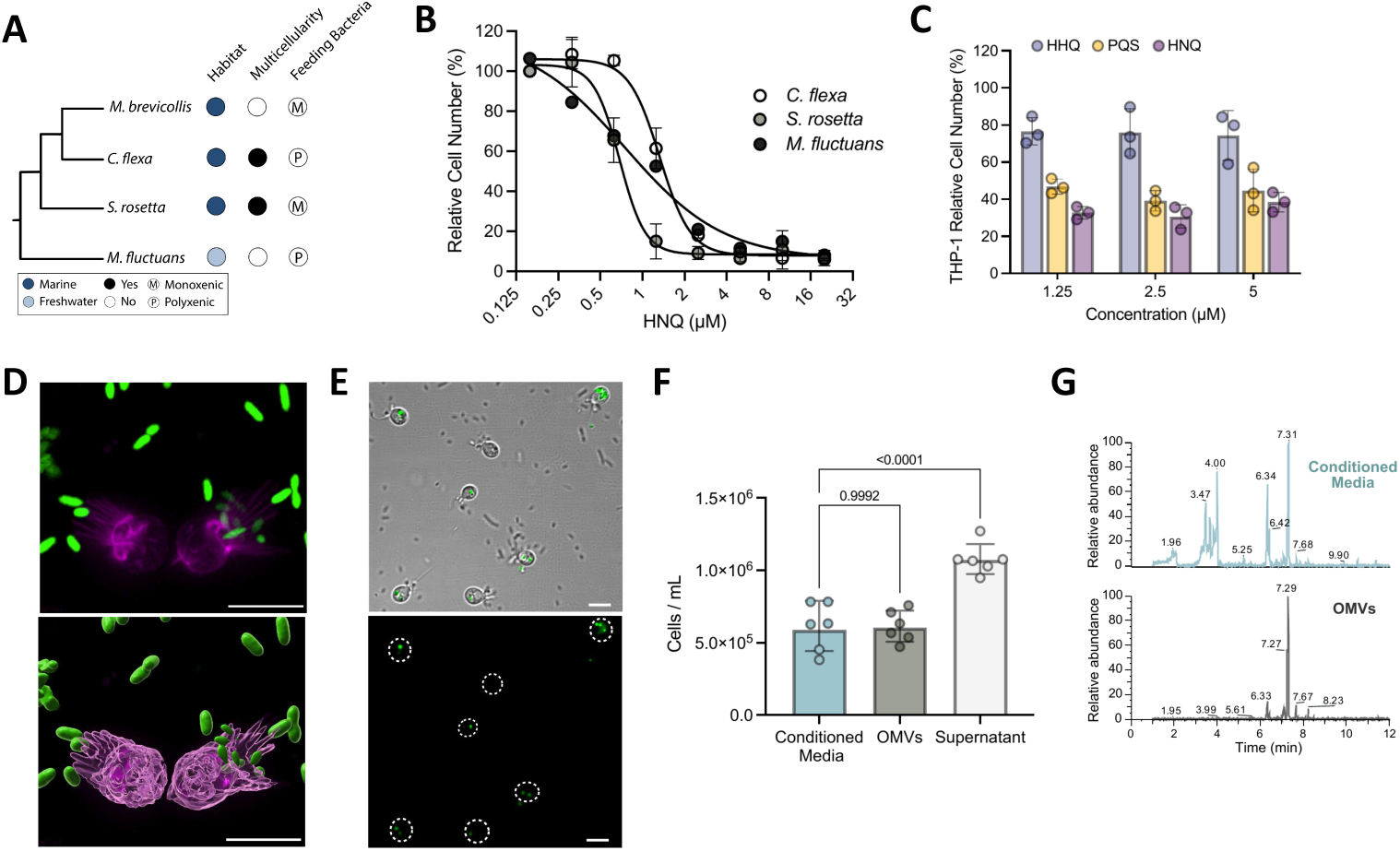
HNQ is a broadly active effector delivered within *P. aeruginosa* outer membrane vesicles **A.** Phylogeny of the four choanoflagellate species tested for susceptibility to *P. aeruginosa* AQs, annotated for habitat (marine or freshwater), multicellularity, and bacterial context (monoxenic or polyxenic culture). **B.** Diverse choanoflagellate species are susceptible to HNQ. Dose-response curves of *C. flexa*, *S. rosetta*, and *M. fluctuans* treated with HNQ; cell counts were obtained after 22 hours (*C. flexa*), 42 hours (*S. rosetta*), or 67 hours (*M. fluctuans*). All three species were sensitive: *C. flexa* (LD50 = 1.33 uM) showed comparable sensitivity to *M. brevicollis* (LD50 = 1.37 uM), while *S. rosetta* (LD50 = 0.68 uM) and *M. fluctuans* (LD50 = 0.75 uM) were approximately twice as sensitive, suggesting HNQ may act on a conserved choanoflagellate target. Data represent mean +/- SD for four biological replicates. **C.** HNQ restricts proliferation of THP-1 human monocytes. THP-1 cells were treated with HHQ, PQS, or HNQ at the indicated concentrations, and relative cell number (%) was quantified after 48 hours. HNQ restricted proliferation at concentrations as low as 1.25 uM and was more potent than PQS, demonstrating activity against human cells at concentrations well below those previously reported for PQS. Data represent mean +/- SD for three biological replicates. **D.** *P. aeruginosa* accumulates at the *M. brevicollis* feeding collar. *M. brevicollis* stably expressing LifeAct-mCherry (magenta, actin) was incubated with GFP-labeled *P. aeruginosa* (PAO1) for three hours and imaged live; panels show maximum-intensity projections (top) and 3D object renderings (bottom). PAO1 is excluded from phagocytosis and instead accumulates at the feeding collar, positioning the bacterium for localized delivery of lipophilic AQs. Scale bars = 10 µm. **E.** OMVs are taken up by *M. brevicollis*. Representative DIC (top) and fluorescence (bottom) images of *M. brevicollis* incubated for 30 minutes with OMVs stained with the lipophilic dye FM 4-64FX (green). Dashed circles mark cell positions; intracellular OMV signal is visible in a subset of cells, indicating that OMVs are internalized. Scale bars = 5µm. **F.** PAO1 OMVs recapitulate the cytotoxicity of conditioned media. *M. brevicollis* was treated with *P. aeruginosa* conditioned media, isolated OMVs, or OMV-depleted supernatant (OMVs removed by ultracentrifugation) at matched equivalents, and cell density was quantified after 22 hours. Each point represents one biological replicate; bars represent mean +/- SD. P-values are the result of an ordinary one-way ANOVA. **G.** AQs are incorporated into outer membrane vesicles (OMVs). UHPLC-HRMS chromatograms comparing *P. aeruginosarhlA^-^* conditioned media and isolated OMVs. Identified quinolones occur at 6.34 or 6.33 min (PQS), 6.42 or 6.41 min (HHQ), and 7.31or 7.29 min (HNQ).

The role of quorum sensing in *P. aeruginosa* pathogenesis has motivated prior investigation of how human cells respond to purified PQS, which has reported antiproliferative or cytotoxic effects at concentrations between 10–100 µM(18–21, 23). Because choanoflagellates are susceptible to HNQ at much lower concentrations than PQS, we asked whether HNQ similarly affects human cell viability. We first assessed the dose-dependent response of THP-1 monocytes to HNQ, HHQ, and PQS. Although none of the AQs were cytotoxic as measured by LDH release (Figure S3C), HNQ restricted cell proliferation at concentrations as low as 1.25 µM and was more potent than PQS (Figure 4C). Examining the response of A549 epithelial airway cells revealed that HNQ and PQS only modestly reduce proliferation at the tested concentrations (Figure S3E), which are lower than those used in many published studies but selected based on HNQ aqueous solubility. Nonetheless, these results demonstrate that HNQ can directly affect human cell viability at biologically relevant concentrations.

### HNQ reaches *M. brevicollis* through cell contact and outer membrane vesicles

Our finding that HNQ reduces the viability of choanoflagellates and human cells raises the question of how this lipophilic molecule reaches target cells in the aquatic environments where choanoflagellates would encounter *P. aerugino*sa, or during infection of a mammalian host. In addition, although choanoflagellates phagocytose most planktonic bacteria as prey, we previously observed that *P. aeruginosa* (PAO1) is not eaten by *M. brevicollis*(2), further motivating us to investigate how HNQ reaches *M. brevicollis*. We incubated *M. brevicollis* with fluorescent *P. aeruginosa* (PAO1) or *E. coli* (BW25113) for 3 hours and visualized choanoflagellate-bacterial interactions using live imaging. Consistent with our previous findings and in contrast to *E. coli*, PAO1 was not detected within phagocytic cups or intracellular food vacuoles (Figure. S4A; Figure 4). Instead, PAO1 accumulated at the *M. brevicollis* feeding collar (Figure 4D). This sustained close contact between *M. brevicollis* and planktonic *P. aeruginosa* could facilitate delivery of lipophilic AQs in aquatic environments.

A second potential delivery route is packaging within outer membrane vesicles (OMVs). PQS and other virulence factors have been shown to partition into *P. aeruginosa* OMVs, which can fuse with host cells to deliver their contents(23, 49–54). We therefore investigated whether OMVs serve as a vehicle for HNQ. OMVs isolated from late stationary-phase cultures recapitulated the cytotoxic effect of conditioned media and were observed intracellularly in *M. brevicollis* (Figure 4E,F). In contrast, OMV-depleted supernatant, from which OMVs had been removed by ultracentrifugation, did not affect *M. brevicollis* viability (Figure 4F). Comparison of the UHPLC-HRMS chromatograms from conditioned media and OMV extracts indicated that multiple AQs are incorporated into OMVs (Figure 4G). We therefore quantified HNQ, HHQ, and PQS in OMV and supernatant extracts. Strikingly, HNQ was highly enriched in OMVs and nearly absent from the supernatant, while HHQ was detected at similar concentrations in both OMVs and supernatant (Table 4). PQS, in contrast, was enriched in supernatant relative to OMVs (Table 4), consistent with strain-specific differences in PQS export(50). Together, these data support a role for HNQ as a bacterial effector and identify OMVs as a potential delivery mechanism for this hydrophobic molecule in aqueous environments.

### A choanoflagellate ABCG transporter mediates the response to HNQ

We previously found that *M. brevicollis* responds to *P. aeruginosa* with widespread transcriptional changes, including the induction of STING signaling (2). To understand how HNQ contributes to this response, we set out to investigate the host pathways it activates. We first tested whether STING signaling mediates responses downstream of HNQ by comparing the dose-dependent survival of wildtype and *STING^-^* cells. Both wildtype and *STING^-^* cells were similarly susceptible to HNQ (Figure S5A), indicating that STING-independent pathways are likely involved in responding to this effector.

We therefore used RNA-seq and differential expression analysis to identify candidate pathways activated in response to HNQ. We treated *M. brevicollis* with either HNQ (1.25 µM) or *P. aeruginosa* conditioned media for three hours, then quantified differential gene expression relative to untreated cells for each condition. HNQ and conditioned media induced distinct transcriptional responses in *M. brevicollis* (Figure 5A; Figure S5B,C). 952 genes were differentially expressed in response to conditioned media treatment (|log_2_FC|>1 and adj. p-value<0.05) (Figure S5C), which is consistent with previous experiments. In contrast, HNQ treatment yielded only 108 significantly differentially expressed genes, though the high prevalence of uncharacterized genes limited our ability to identify additional biological pathways involved.

**Figure 5.**
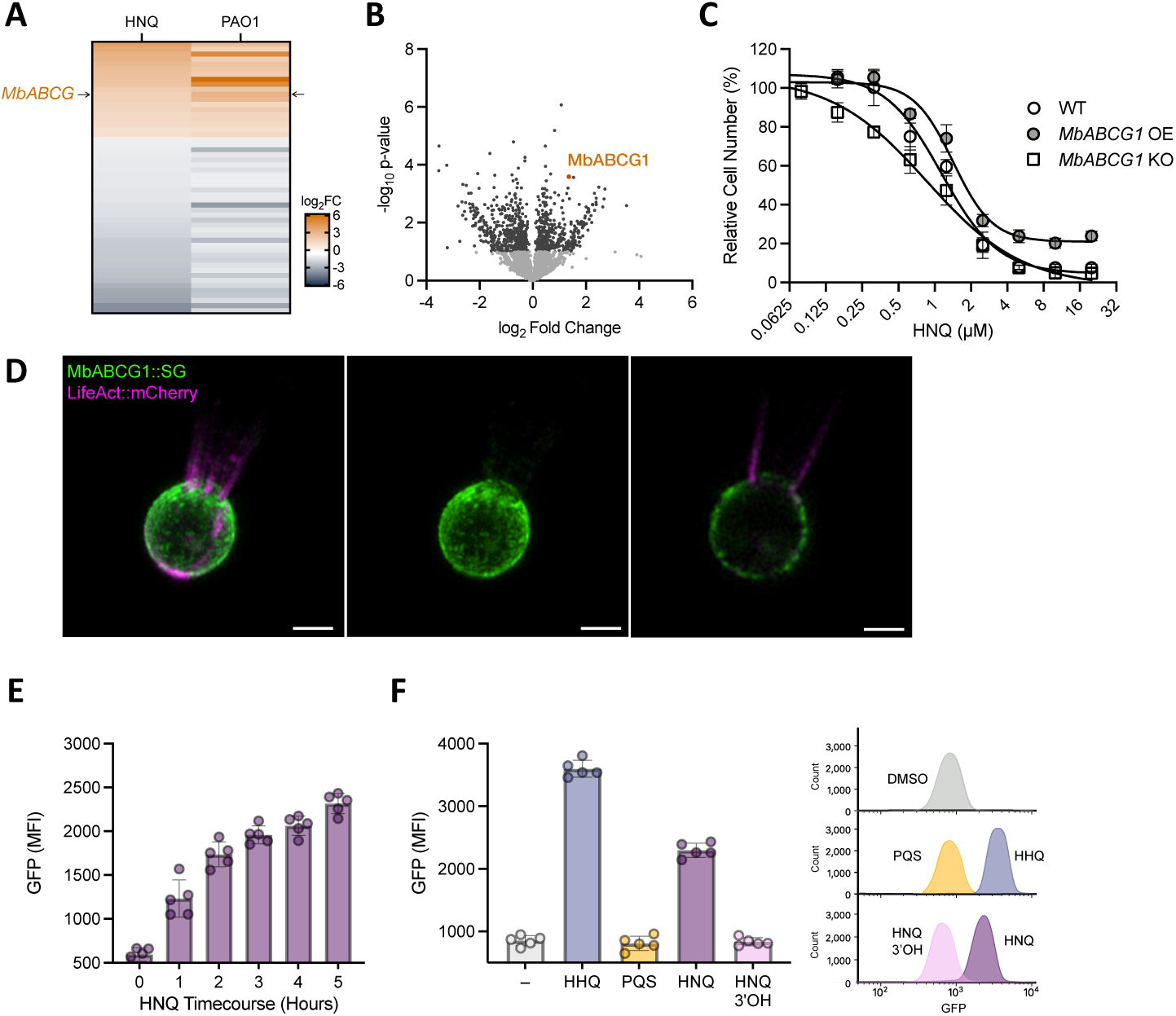
Novel choanoflagellate transporter MbABCG1 responds selectively to non-hydroxylated alkylquinolones and protects against HNQ. **A.** HNQ and *P. aeruginosa* conditioned media induce distinct but overlapping transcriptional responses in *M. brevicollis*. Heatmap comparing the log_2_ fold-change expression of HNQ-responsive genes that are significantly differentially expressed, across HNQ and PAO1 conditioned media treated cells. **B.** Volcano plot depicting the relative protein abundance in *M. brevicollis* treated with 1.25 µM HNQ compared to untreated control cells. PSMs for three biological replicates were generated in Proteome Discover to enable Degust usage. Differential protein enrichment was computed using the RUVr edeR-quasi-likelihood model implement in Degust, requiring a minimum count of two PSMs in at least two samples, and normalizing by Trimmed Mean of M-values. **C.** MbABCG1 expression modulates *M. brevicollis* sensitivity to HNQ. Dose-response curves comparing wild-type (WT) cells to MbABCG1 knockout (*MbABCG1^-^)* (D) and overexpression (*MbABCG1* OE) (E) cell lines treated with HNQ. Loss of *MbABCG1* sensitizes cells to lower concentrations of HNQ. Overexpression of *MbABCG1* promotes survival in the presence of high concentrations of HNQ. Relative cell number (%) was calculated for each genotype against its own DMSO control. Data represent mean +/- SD for four biological replicates. **D.** MbABCG1 is localized to the plasma membrane. Representative maximum intensity projection of the endogenously tagged MbABCG1::StayGold. Co-expression a *LifeAct:mCherry* reporter construct (magenta) indicates the apical collar and orients apical-basal axis of cell. MbABCG1::SG (green) is punctate but distributed uniformly at the plasma membrane, as further highlighted by Z-slice (right). Scale bars = 2 µm. **E, F.** MbABCG1 abundance, measured by MbABCG1::SG median fluorescence intensity (MFI), increases in response to HNQ and non-hydroxylated alkylquinolones, but not hydroxylated congeners. **(E)** Flow cytometry time course of MbABCG1::SG MFI following treatment with 1.25 µM HNQ. **(F)** (Left) Cells were treated with a DMSO vehicle control, or 1.25 µM of the indicated AQs for five hours and analyzed by flow cytometry. Non-hydroxylated AQs markedly increased MbABCG1 abundance compared to DMSO control, whereas hydroxylated congeners did not. Each point on bar graphs represents one biological replicate; bars represent mean +/- SD for 5 biological replicates. (Right) Flow cytometry histograms of show a distinct, population-wide shift in fluorescence following treatment with HHQ and HNQ, but not their 3’ hydroxylated analogs. MbABCG1 cell count and fluorescence is the average of 5 biological replicates.

To prioritize candidates for further study, we examined the overlap of genes differentially expressed in response to HNQ and conditioned media. This analysis identified 54 genes that were significantly differentially expressed (19 upregulated, 35 downregulated) in response to both stimuli. Shotgun proteomics confirmed that *MONBRDRAFT_34196*, a predicted ABCG half transporter that we initially identified through RNA-seq, is also upregulated at the protein level following HNQ treatment (Figure 5B). Therefore, we decided to further explore the function of *MONBRDRAFT_34196*, hereafter referred to as *MbABCG1,* to investigate the cellular response to HNQ and parse specific pathways involved in the response to *P. aeruginosa*.

*MbABCG1* is predicted to encode an ABCG half-transporter, comprising a single nucleotide-binding domain (NBD) and one six-pass transmembrane domain (Figure S5D). Phylogenetic analysis of MbABCG1 placed it within a choanoflagellate-specific clade, separate from functionally characterized animal ABCG subfamilies (Figure S6). This lineage-specific placement makes the substrates and function of MbABCG1 enigmatic.

To test whether MbABCG1 mediates the *M. brevicollis* response to HNQ, we generated loss- and gain-of-function cell lines. Using CRISPR-mediated genome editing, we disrupted *MbABCG1* within the first exon of its coding region, targeting the nucleotide-binding domain (*MbABCG1*^-^),and in parallel, generated a transgenic line that stably overexpresses the *MbABCG1* coding sequence under the actin promoter (*MbABCG1* OE). Neither manipulation altered baseline proliferation, as all three cell lines grew comparably in the absence of HNQ (Figure S5E). We next compared the viability of wildtype, *MbABCG1*^-^, and *MbABCG1* OE cells across a range of HNQ concentrations. Overexpression of *MbABCG1* promoted survival, both at high HNQ concentrations that were lethal to wildtype cells (Figure 5C), and notably, when challenged with PAO1 conditioned media (Figure S5F). *MbABCG1*^-^ cells were more susceptible to HNQ than wildtype cells, losing viability at concentrations that did not appreciably affect wildtype survival (Figure 5C).

To examine the subcellular localization and dynamics of MbABCG1, we established a protocol for generating endogenously tagged fusion proteins in *M. brevicollis*. We engineered a knock-in cell line in which a short linker followed by a monomeric StayGold fluorescent protein was inserted immediately before the *MbABCG1* stop codon. This tag placement was guided by a prior study which determined that a C-terminal fluorescent protein does not impair ABCG transporter function^37^. Confocal microscopy revealed that MbABCG1 localizes to the plasma membrane, where its distribution appears uniform (Figure 5D).

Although MbABCG1::SG was detectable across the population at steady state, widefield imaging suggested that HNQ treatment increased the fluorescence of individual cells. To quantify this response, we used flow cytometry to measure MbABCG1::SG median fluorescence intensity (MFI) and confirmed that MbABCG1 abundance increases following HNQ exposure (Figure 5E). We found that this upregulation was specific to non-hydroxylated AQs: HHQ (C_7_), 2-octyl-4-quinolone (C_8_), and HNQ (C_9_) each increased MbABCG1 abundance, whereas the hydroxylated congeners PQS and 3’-hydroxy-HNQ did not, even when tested at higher concentrations (Figure 5F; Figure S5H). Cyclic dinucleotides (2’3’- and 3’3’-cGAMP), which elicit broad transcriptional responses in *M. brevicollis*, also failed to increase MbABCG1 (Figure. S5I), indicating that upregulation is not a generic consequence of immune activation. Finally, conditioned media from wild-type, but not AQ-deficient (*pqsA*), *P. aeruginosa* produced a subtle but reproducible increase in MbABCG1 abundance (Figure S5J), linking this response to bacterially secreted AQs. Together, these results show that MbABCG1 abundance increases specifically in response to HNQ and other non-hydroxylated AQs. This selectivity mirrors our earlier finding that a sublethal dose of HNQ enhances the potency of HHQ but not PQS (Figures S2B,C), providing a separate line of evidence that *M. brevicollis* distinguishes non-hydroxylated from hydroxylated AQs.

## Discussion

Here, we identify HNQ, an abundant but overlooked *P. aeruginosa* alkylquinolone, as a potent effector that reduces the viability of choanoflagellates and human cells, and we define the chemical features required for its bioactivity in the choanoflagellate *M. brevicollis*. Using HNQ as a defined agonist, we show that *M. brevicollis* mounts distinct responses to non-hydroxylated and hydroxylated AQs, and uncover a novel choanoflagellate ABCG transporter, MbABCG1, that specifically mediates responses to HNQ and promotes survival. Together, these findings expand the set of *P. aeruginosa* effectors relevant to host-pathogen interactions, further our understanding of how choanoflagellates recognize bacterial pathogens, and provide a new probe for dissecting choanoflagellate immune response pathways.

Choanoflagellates inhabit aquatic environments where they simultaneously capture bacteria as prey and contend with bacterial pathogens. Successfully navigating these interkingdom interactions requires choanoflagellates to precisely interpret a complex milieu of bacterially produced chemical signals. Previous work has shown the stringency with which choanoflagellates respond to bacteria in developmental contexts(5–7, 55–57); for example, minor structural differences between bacterial sulfonolipids markedly alter their ability to induce *S. rosetta* multicellular development(5, 55). Our findings that *M. brevicollis* discriminates among closely related AQs, mounting distinct responses to those differing by a single hydroxyl group, extend this theme to host-pathogen interactions. Because some AQs, including HNQ, are produced by a select few bacterial species, these molecules could serve as chemical signatures that help choanoflagellates identify a specific pathogen within a broad bacterial community.

The interkingdom signaling potential of *P. aeruginosa* alkylquinolones has been studied primarily through the quorum-sensing molecules PQS and HHQ. Nonetheless, HNQ was initially isolated over 80 years ago as Pyo Ic in a search for antibiotics(28, 58) and is among the more abundant alkylquinolones produced by *P. aeruginosa*(29). Why then, has HNQ not been previously implicated in interkingdom interactions or pathogenesis? It is possible that HNQ has been largely ignored because it showed little antibacterial activity following its discovery and, in more recent years, has been overshadowed by PQS and HHQ, whose roles in quorum sensing have made them the primary subjects of host–AQ studies. In addition, because many AQs share a common biosynthetic pathway(39), the individual contributions of each congener are difficult to disentangle genetically during host–pathogen interactions. Our unbiased, activity-guided approach allowed us to resolve HNQ as a discrete effector that is bioactive towards diverse choanoflagellates and human cells, and similar methods may prove powerful for uncovering the roles of other understudied small molecules.

Systematically testing HNQ and its congeners allowed us to define the structural features that confer activity in choanoflagellates. Shortening or lengthening the alkyl chain of HNQ, or adding a 3′-hydroxyl group, reduces both potency and efficacy, suggesting that HNQ acts through a specific interaction with a host protein rather than through nonspecific, detergent-like effects. Our data also indicate that non-hydroxylated (HNQ, HHQ) and hydroxylated (PQS) AQs engage distinct *M. brevicollis* pathways: a sublethal concentration of HNQ additively enhanced the potency of HHQ but not PQS, and MbABCG1 abundance increased specifically in response to non-hydroxylated AQs but not their 3′-hydroxylated counterparts. Although the mechanisms by which AQs affect host biology remain largely unresolved, our findings parallel other eukaryotic systems, wherein HHQ and PQS elicit different responses. For example, HHQ but not PQS arrests the growth of diatoms and coccolithophores at physiologically relevant concentrations(22, 24), PQS but not HHQ destabilizes HIF-1α in airway epithelial cells(19), and although both PQS and HHQ suppress NF-κB–mediated gene expression in mouse macrophages, they likely do so by different mechanisms(18).

Alkylquinolones are lipophilic molecules with low aqueous solubility, raising the question of how they reach target cells in aquatic environments. We found that *P. aeruginosa* outer membrane vesicles are internalized by *M. brevicollis* and sufficient to reduce viability, and that HNQ is enriched in OMVs. Thus, packaging within OMVs provides a plausible mechanism for concentrating and delivering HNQ in aqueous environments, analogous to the OMV-mediated transport of PQS described in other systems(49, 51, 53, 54). While OMV biogenesis and composition undoubtedly vary in response to specific environmental conditions, the partitioning of HNQ into OMVs reported here suggests that this molecule could serve as an effector in a wide range of contexts, including in mammalian infection.

We identified a novel choanoflagellate ABCG half-transporter, MbABCG1, that mediates a protective response to *P. aeruginosa* alkylquinolones. Developing a protocol to endogenously tag *MbABCG1* allowed us to observe its localization at the plasma membrane, and more importantly, discern that its abundance increases in response to non-hydroxylated, but not hydroxylated, AQs. This upregulation also occurs with conditioned media isolated from wild-type, but not AQ-deficient (*pqsA*), *P. aeruginosa*, indicating that MbABCG1 responds to even low concentrations of AQs within a complex secretome. Consistent with a protective role, overexpression of MbABCG1 promotes survival following exposure to HNQ or *P. aeruginosa* conditioned media, whereas its deletion increases sensitivity to HNQ. Although its substrates remain unknown, MbABCG1 belongs to a clade of choanoflagellate-specific transporters; because HNQ is similarly potent in other species, related transporters may mediate protective responses across choanoflagellates. Whether MbABCG1 directly exports or sequesters AQs, or instead transports host-derived molecules involved in the response to these effectors, remains an important question for understanding the mechanism of this protection.

Finally, few defined bacterial effectors are available for dissecting host–pathogen interactions in choanoflagellates. Whereas previous studies relied on *P. aeruginosa* live bacteria or conditioned media, which elicit hundreds of transcriptional changes, HNQ provides a new agonist for investigating host responses across choanoflagellate species. Its activity also extends to human cells, where it restricts proliferation at concentrations below those reported for PQS, raising the possibility that HNQ contributes to *P. aeruginosa* pathogenesis in mammals. More broadly, using HNQ as a probe across different hosts and contexts may identify previously unrecognized roles for alkylquinolones in interkingdom signaling, from environmental interactions to infection.

## Supporting information

Key Resource Details

Physical data and NMR spectra

## ACKNOWLEDGEMENTS

We thank the Woznica lab for valuable feedback throughout the project, and Kirstin Rollins for assistance with flow cytometry. We thank Despoina Mavridou, Bryan Davies, and Jaime Lopez for sharing bacterial strains and mammalian cell lines. We thank Thomas Böttcher for helpful discussion related to quinolone signaling. We dedicate this work to the memory of Jon Clardy, whose mentorship and generosity brought our groups together and made this collaboration possible.

Super-resolution microscopy and flow cytometry were performed with the support of the UT Austin Microscopy and Flow Cytometry Facility (RRID:SCR_021756). We greatly acknowledge the ERC Starting Grant project 802736 MORPHEUS for funding.

This work was supported by a Hanna H. Gray Fellowship from the Howard Hughes Medical Institute (A.W.) and by the National Institute of General Medical Sciences of the National Institutes of Health under Award Number R35GM165755 (A.W.). The content is solely the responsibility of the authors and does not necessarily represent the official views of the National Institutes of Health.

## Author contributions

A.W. and C.B. designed research; J.G., C.M., M.D., C.P., V.D., F.-X.M., and A.W. performed research; J.G., C.M., M.D., C.P., V.D., F.-X.M., C.B., and A.W. analyzed data; J.G., C.M., M.D., C.B. and A.W. wrote the paper.

---

**Figure S1.** Bioactive fractions 12 and 14 contain distinct alkylquinolone congeners **A.** Parental and rhamnolipid-deficient *P. aeruginosa* strains have similar secreted metabolomic profiles. UHPLC-HRMS total ion chromatograms of organic extracts from the parental mPAO1 strain (top) and the rhamnolipid-deficient *rhlA::Tn* strain (bottom) conditioned media. Numbers above peaks indicate retention times (min). Extracts were analyzed at 50 ug/mL in positive ionization mode. **B.** Bioactive fractions 12 and 14 each contain distinct alkylquinolone congeners. UHPLC-HRMS chromatograms of the *P. aeruginosarhlA::Tn* crude organic extract (top), Fraction 12 (middle), and Fraction 14 (bottom). Fraction 12 contains predominate peaks at 6.33 min and 6.42 min, while Fraction 14 contains a predominate peak at 7.29 min.

**Figure S2.** Synergistic effects are not observed between HNQ and other *P. aeruginosa* alkylquinolones or diketopiperazines. **A.** Additional structure-activity analysis of HNQ congeners. Dose-response curves of *M. brevicollis* treated with HNQ (3) or compounds 5, 10, and 11 (structures shown in Fig. 2A). **B.** Co-treatment with HNQ additively enhances the potency of HHQ. *M. brevicollis* was treated with HNQ alone, HHQ alone, or HHQ titrated in the presence of a fixed sublethal concentration of HNQ (0.625 uM). **C.** Co-treatment with HNQ does not affect the potency or efficacy of PQS. *M. brevicollis* was treated with HNQ alone, PQS alone, or PQS titrated in the presence of 0.625 uM HNQ. Unlike HHQ, the PQS curve is unchanged by HNQ, raising the possibility that hydroxylated (PQS) and non-hydroxylated (HHQ, HNQ) alkylquinolones are perceived through distinct *M. brevicollis* receptors. **D.** AQ-deficient *P. aeruginosa* still restricts *M. brevicollis* growth relative to non-pathogenic bacteria. *M. brevicollis* was treated with conditioned media from a non-pathogenic control bacterium (*Pseudoalteromonas shioyasakiensis*), parental PAO1 strain (WT; mPAO1), or the *pqsA*, *pqsD*, *pqsB* or *pqsH* mutant strains. AQ-null mutants suppressed growth more than the *Pseudoalteromonas* control, indicating that additional secreted factors contribute to cytotoxicity. Each point represents one biological replicate; bars represent mean +/- SD. Statistical analysis was performed using a RM one-way ANOVA. Shown are the adjusted P-values from a Dunnett’s multiple comparison test. **E.** Purified *P. aeruginosa* diketopiperazines (DKPs) do not reduce *M. brevicollis* viability. *M. brevicollis* was treated with Cyclo(Phe-Pro) or Cyclo(Tyr-Pro) — cyclic dipeptides isolated from inactive HPLC fractions 2-7 — across the indicated concentrations. Neither DKP reduced viability. **F.** DKPs do not act synergistically with HNQ. *M. brevicollis* was treated with a sublethal (0.625 µM) or intermediate (1.25 µM) concentration of HNQ, alone or in combination with 20 µM DKPs (10 µM each of Cyclo(Phe-Pro) and Cyclo(Tyr-Pro)). Each point represents one biological replicate; bars represent mean +/- SD. An ordinary one-way ANOVA was used to test for statistical significance; ns, not significant. For all experiments, *M. brevicollis* viability was evaluated after 22 hours, and relative cell number (%) was calculated against a DMSO control. Data represent mean +/- SD for at least three biological replicates.

**Figure S3.** Diverse choanoflagellates and human cells are susceptible to HNQ, an effector primarily produced by *P. aeruginosa* **A.** Structural requirements for AQ activity are conserved across choanoflagellates. Dose-response curves of *C. flexa*, *S. rosetta*, and *M. fluctuans* treated with HHQ, for direct comparison with the HNQ curves in Figure 4B. HNQ was 5-7 fold more potent than HHQ across all species tested, indicating that the preference for the C9 side chain is conserved. Data represent mean +/- SD for at least three biological replicates. **B.** HNQ and related congeners reduce the viability of *S. rosetta* single cells and multicellular rosettes. Uninduced (single cells) and rosette-induced (multicellular) *S. rosetta* cultures were treated with the indicated concentrations of Fraction 14. Relative cell number was calculated with respect to a DMSO control. Data represent mean +/- SD for two biological replicates. **C.** AQs do not cause acute membrane lysis in THP-1 cells. Lactate dehydrogenase (LDH) release (RFU) from THP-1 cells treated with HHQ, PQS, or HNQ at the indicated concentrations, with DMSO as a negative control and DMSO + Triton X-100 as a lysis-positive control. AQs did not increase LDH release above the DMSO control after 48 hours. Data represent mean +/- SD for three biological replicates. **D.** Absolute THP-1 cell density (cells/mL) following treatment with DMSO, HHQ, PQS, or HNQ at the indicated concentrations, quantified after 48 hours. HNQ and PQS significantly reduced cell number relative to DMSO at all concentrations tested, whereas HHQ had lesser effects. Each point represents one biological replicate; bars represent mean +/- SD. Shown are multiplicity adjusted P-values (Dunnett) following a two-way ANOVA test. **E.** HNQ only modestly affects the proliferation of A549 human airway epithelial cells. A549 cells were treated with HHQ, PQS, or HNQ at the indicated concentrations and relative cell number (%) was quantified after 24 hours relative to a DMSO control. Data represent mean +/- SD for four biological replicates. **F, G.** The secreted metabolomes of diverse *Pseudomonas* species are distinct. (F) Principal component analysis (PCA) of the secreted metabolomes of *P. aeruginosa* PAO1, ARB769, and five other *Pseudomonas* species reveals clear species-level separation. Axes indicate the proportion of variance. (G) Hierarchical clustering of the same secreted metabolomes confirms intra-species reproducibility and groups the two *P. aeruginosa* strains (PAO1, ARB769) apart from the other species.

**Figure S4.** *M. brevicollis* internalizes *E. coli* but not *P. aeruginosa* PAO1. **A.** *M. brevicollis* stably expressing LifeAct-mCherry (magenta, actin) was incubated with GFP-labeled *E. coli* for three hours and imaged live; panels show maximum-intensity projections (top) and 3D object renderings (bottom) highlighting bacteria in phagocytic cups and intracellular food vacuoles. Scale bars = 10 µm.

**Figure S5.** MbABCG1 is involved in responding to HNQ and non-hydroxylated AQs **A.** STING signaling does not affect survival following HNQ exposure. Dose-response curves of wild-type (WT) and STING knockout (STING KO) *M. brevicollis* cell lines treated with HNQ. Data represent mean +/- SD for three biological replicates. **B.** Transcriptome-wide response to HNQ and *P. aeruginosa* conditioned media. Heatmap of log_2_ foldchange for all genes following a three hour exposure to *P. aeruginosa* conditioned media (left) or HNQ (right), relative to untreated cells. **C.** Venn diagram of genes significantly differentially expressed in *M. brevicollis* after 3 hours of treatment with HNQ (1.25 µM) or PAO1 conditioned media, relative to untreated cells (|log2FC| > 1, adjusted p < 0.05). **D.** MbABCG1 (*MONBRDRAFT_34196*), identified among the HNQ-responsive genes and confirmed to be upregulated at the protein level by shotgun proteomics, is predicted to be a multi-pass transmembrane ABCG-family transporter. Schematic of the MbABCG1 half-transporter protein domain architecture, showing a cytoplasmic N-terminal nucleotide-binding domain (NBD) followed by a six-pass transmembrane domain Scissors indicate the CRISPR/Cas9 editing site used to generate knockout cell line. SG indicates placement of StayGold protein sequence at C-terminus. **E.** *MbABCG1* knockout or overexpression does not affect cell growth rates. *M. brevicollis* wildtype, *ABCG1-* and *pActin::ABCG1* cell lines were seeded at the same cell density and grown for 24 hours under standard culturing conditions. Data represent mean +/- SD for four biological replicates. **F.** MbABCG1 expression modulates *M. brevicollis* sensitivity to PAO1 conditioned media. Wild-type (WT), MbABCG1 overexpression (*MbABCG1* OE), and MbABCG1 knockout (*MbABCG1^-^)* cell lines were treated with conditioned media isolated from *P. aeruginosa* wild-type (mPAO1) or AQ-deficient (*pqsA*) strains. Overexpression of *MbABCG1* promotes survival in the presence of mPAO1. *MbABCG1* knockout cells are more sensitive to mPAO1 than *pqsA* conditioned media, as observed for parental *M. brevicollis.* Data represent mean +/- SD for three biological replicates. **G.** Flow cytometry quantification of MbABCG1::SG MFI (GFP channel) in the endogenously tagged MbABCG1::SG knock-in line compared with untagged parental *M. brevicollis*. **H.** MbABCG1 abundance increases in response to non-hydroxylated alkylquinolones, but not hydroxylated congeners. Flow cytometry quantification of MbABCG1::SG median fluorescence intensity (GFP) after 5-hour treatment 1.25 µM non-hydroxylated AQs, and 1.25 or 5 µM hydroxylated AQs. **I.** MbABCG1 abundance does not change in response to endogenous or bacterially-produced cyclic dinucleotides. Cells were treated with a water vehicle control, or 100 µM 2’3’ cGAMP or 3’3’ cGAMP for five hours and analyzed by flow cytometry. Each point on bar graph represents one biological replicate; bars represent mean +/- SD for 5 biological replicates. **J.** Conditioned media from AQ-producing *P. aeruginosa* increases MbABCG1 abundance. (Left) Flow cytometry quantification of MbABCG1::SG MFI following a 5-hour treatment with an LB control, conditioned media from wild-type *P. aeruginosa* (mPAO1), or conditioned media from the AQ-null *pqsA* mutant (each at 5% vol/vol). WT conditioned media produced a subtle but reproducible increase in MbABCG1 abundance, whereas AQ-deficient *pqsA* conditioned media did not. Each point represents one biological replicate; bars represent mean +/- SD for 5 biological replicates. (Right) Flow cytometry histograms of MbABCG1::SG fluorescence for control (top) and WT or *pqsA* conditioned media-treated (bottom). MbABCG1 cell count and fluorescence represents the average of 5 biological replicates.

**Figure S6.** Phylogenetic placement of MbABCG1 within the opisthokont ABCG transporter family **A.** MbABCG1 belongs to a choanoflagellate-specific clade of ABCG transporters. Pictured is the clade containing MbABCG1, derived from the larger maximum-likelihood tree of opisthokont ABCG transporters built from 335 ABCG sequences representing approximately 100 taxa. **B.** Maximum-likelihood phylogenetic tree of ABCG-family transporters across opisthokonts, with the clade containing MbABCG1 indicated. Clades are collapsed, and color-coding represents the proportion of sequences from different taxonomic groups. The positions of well-studied ABCG transporters, including human ABCG2, ABCG5, ABCG8, scarlet, and white are indicated.

## Notes

### Competing Interest Statement

The authors have declared no competing interest.

