## Supplemental Figures for "Choanoflagellates discriminate among *Pseudomonas aeruginosa* alkylquinolones during pathogen response"

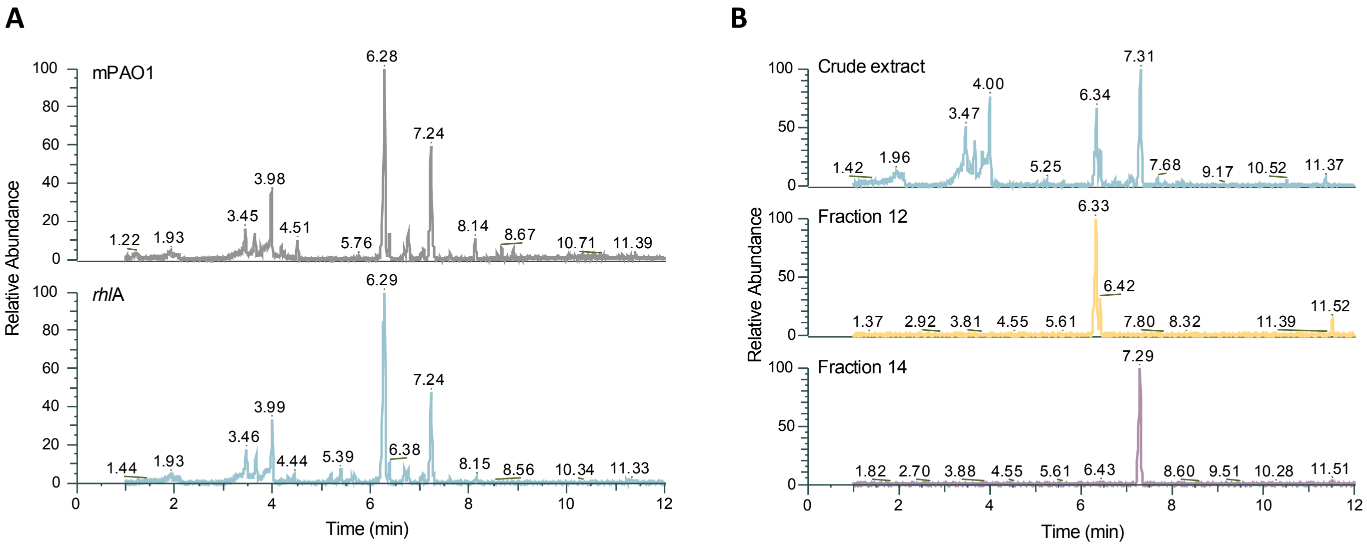
A
mPAO1
rhlA
Relative Abundance
Time (min)
B
Crude extract
Fraction 12
Fraction 14
Relative Abundance
Time (min)

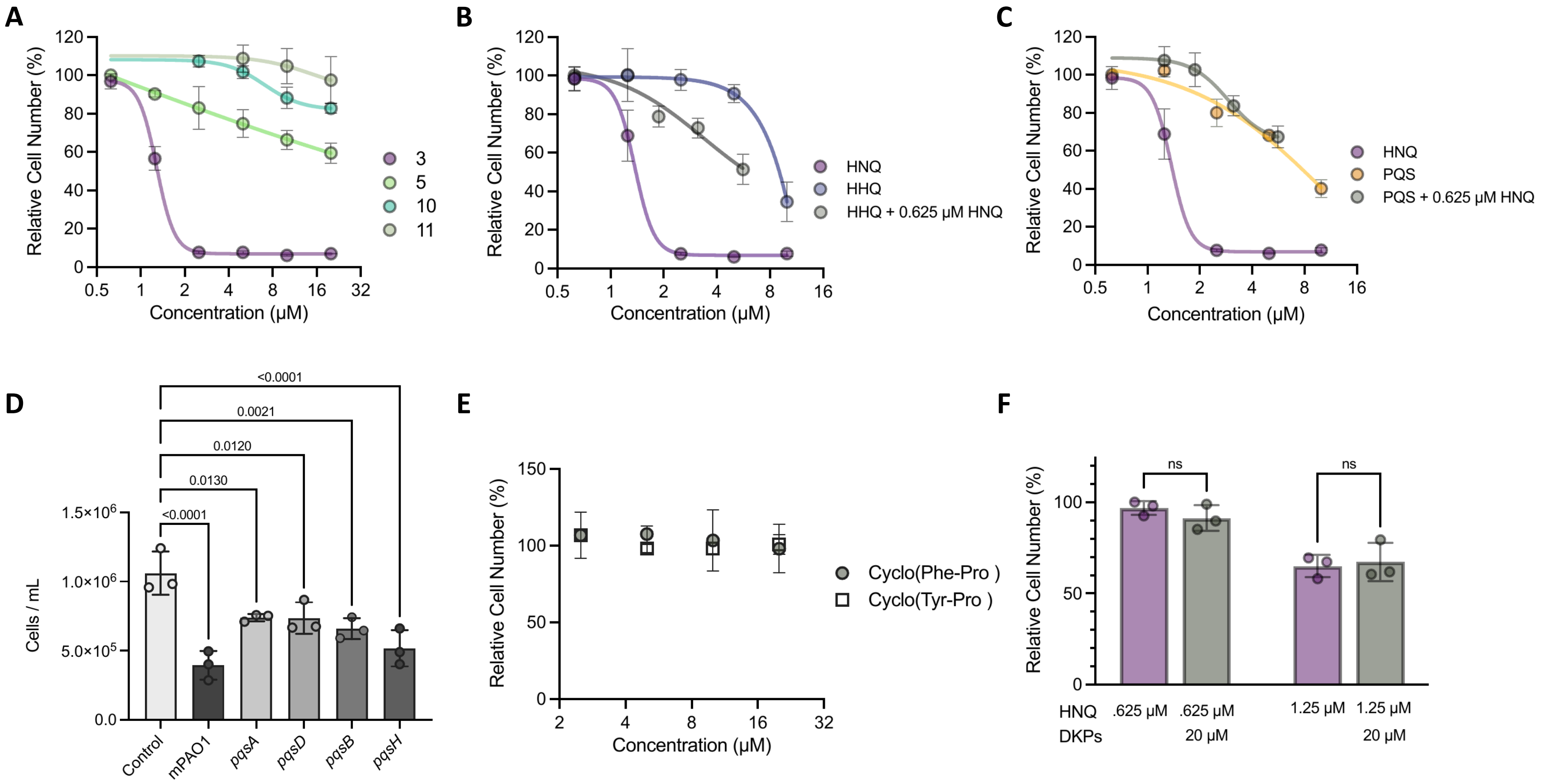

A
Relative Cell Number (%)
Concentration (μM)
3
5
10
11
B
HNQ
HHQ
HHQ + 0.625 μM HNQ
C
HNQ
PQS
PQS + 0.625 μM HNQ
D
Cells / mL
<0.0001
0.0021
0.0120
0.0130
<0.0001
Control
mPAO1
pqsA
pqsD
pqsB
pqsH
E
Cyclo(Phe-Pro )
Cyclo(Tyr-Pro )
F
ns
ns
HNQ .625 μM .625 μM 1.25 μM 1.25 μM
DKPs 20 μM 20 μM

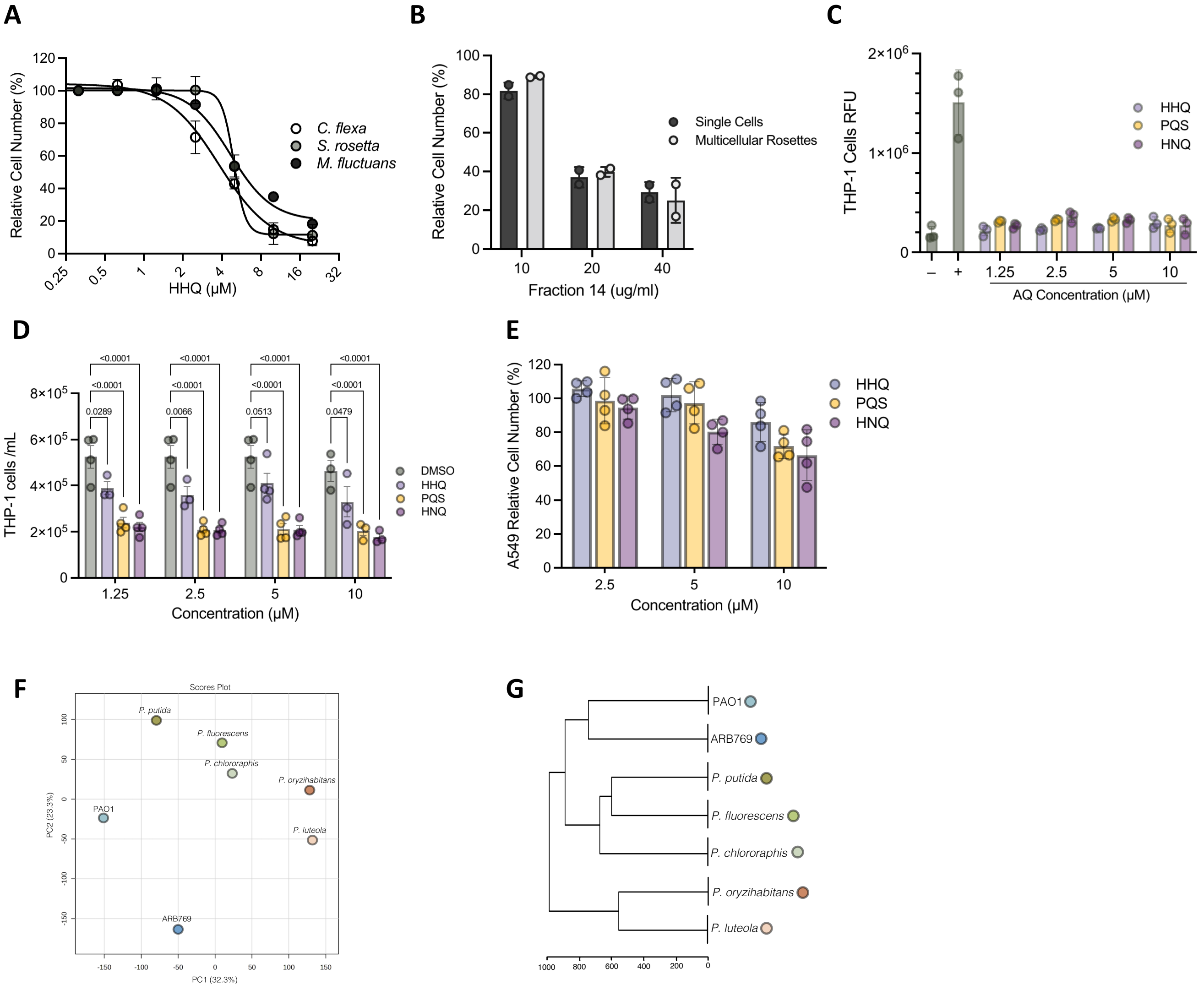

A
Relative Cell Number (%)
HHQ (µM)
C. flexa
S. rosetta
M. fluctuans
B
Relative Cell Number (%)
Fraction 14 (ug/ml)
Single Cells
Multicellular Rosettes
C
THP-1 Cells RFU
AQ Concentration (µM)
HHQ
PQS
HNQ
D
THP-1 cells /mL
Concentration (µM)
DMSO
HHQ
PQS
HNQ
<0.0001
0.0289
0.0066
0.0513
0.0479
E
A549 Relative Cell Number (%)
Concentration (µM)
HHQ
PQS
HNQ
F
Scores Plot
PC1 (32.3%)
PC2 (23.3%)
P. putida
P. fluorescens
P. chlororaphis
P. oryzihabitans
P. luteola
PAO1
ARB769
G
PAO1
ARB769
P. putida
P. fluorescens
P. chlororaphis
P. oryzihabitans
P. luteola

A

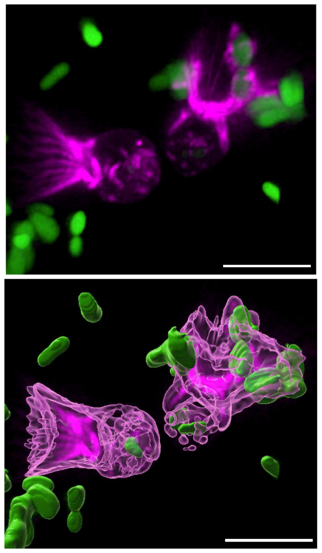

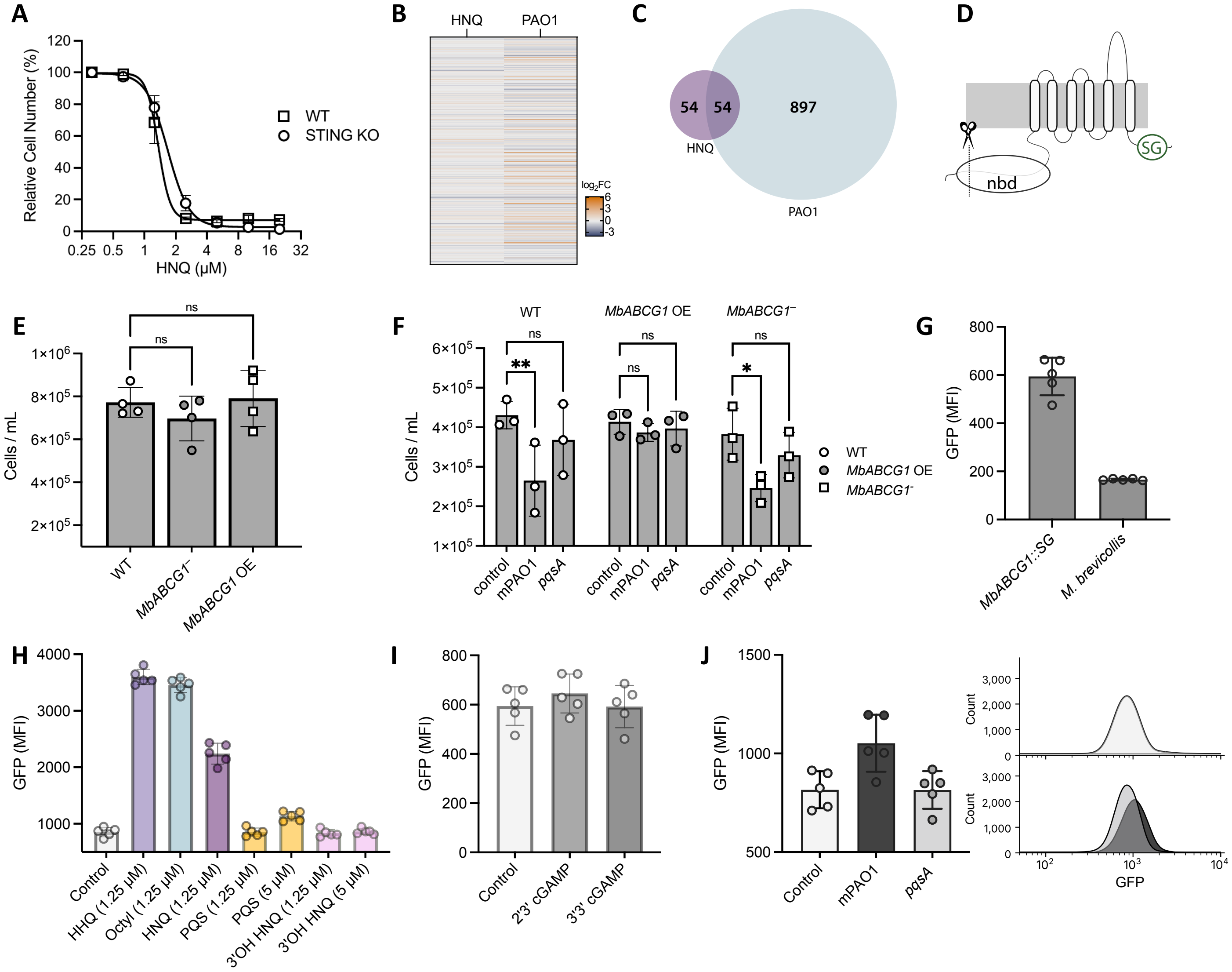
A
Relative Cell Number (%)
HNQ (μM)
WT
STING KO
B
HNQ
PAO1
log2FC
C
54
54
897
HNQ
PAO1
D
nbd
SG
E
Cells / mL
ns
WT
MbABCG1⁻
MbABCG1 OE
F
WT
MbABCG1 OE
MbABCG1⁻
Cells / mL
**
*
ns
control
mPAO1
pqsA
G
GFP (MFI)
MbABCG1::SG
M. brevicollis
H
GFP (MFI)
Control
HHQ (1.25 μM)
Octyl (1.25 μM)
HNQ (1.25 μM)
PQS (1.25 μM)
PQS (5 μM)
3'OH HNQ (1.25 μM)
3'OH HNQ (5 μM)
I
GFP (MFI)
Control
2'3' cGAMP
3'3' cGAMP
J
GFP (MFI)
Control
mPAO1
pqsA
Count
GFP

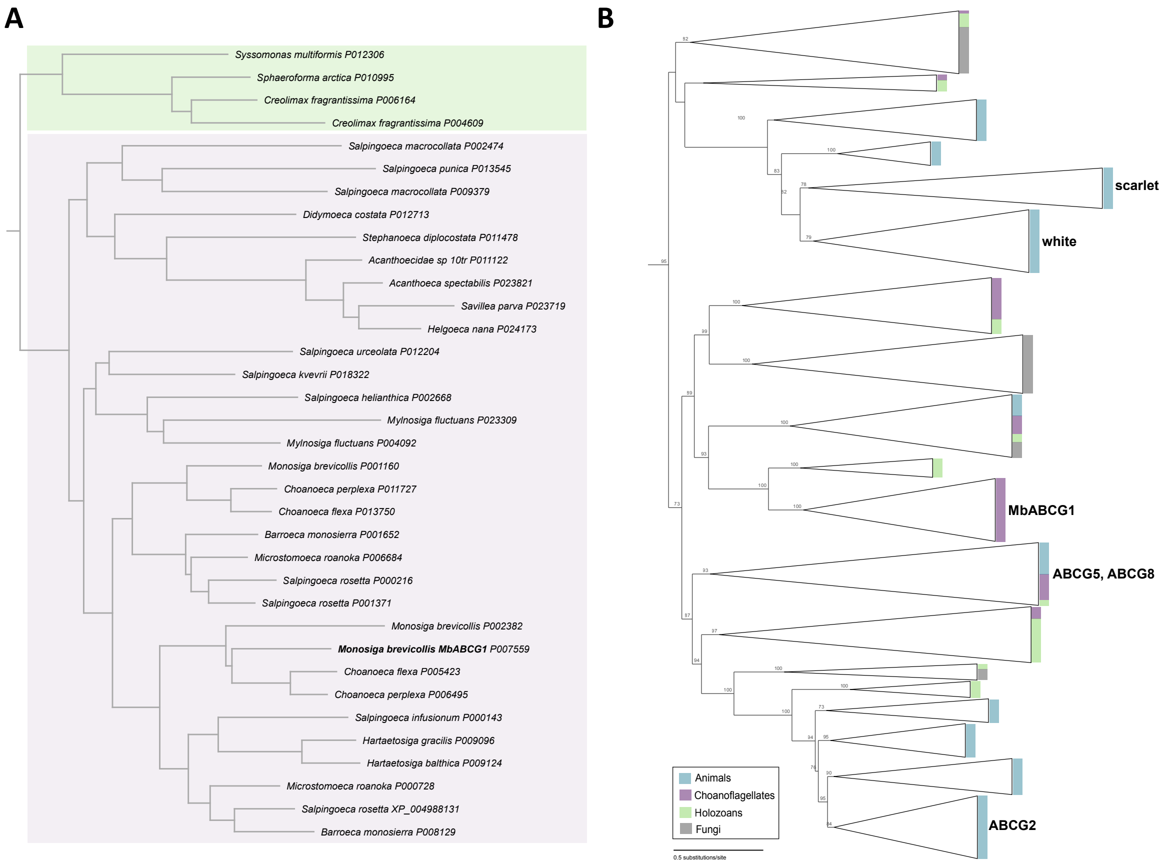
A
Syssomonas multiformis P012306
Sphaeroforma arctica P010995
Creolimax fragrantissima P006164
Creolimax fragrantissima P004609
Salpingoeca macrocollata P002474
Salpingoeca punica P013545
Salpingoeca macrocollata P009379
Didymoeca costata P012713
Stephanoeca diplocostata P011478
Acanthoecidae sp 10tr P011122
Acanthoeca spectabilis P023821
Savillea parva P023719
Helgoeca nana P024173
Salpingoeca urceolata P012204
Salpingoeca kvevrii P018322
Salpingoeca helianthica P002668
Mylnosiga fluctuans P023309
Mylnosiga fluctuans P004092
Monosiga brevicollis P001160
Choanoeca perplexa P011727
Choanoeca flexa P013750
Barroeca monosierra P001652
Microstomoeca roanoka P006684
Salpingoeca rosetta P000216
Salpingoeca rosetta P001371
Monosiga brevicollis P002382
Monosiga brevicollis MbABCG1 P007559
Choanoeca flexa P005423
Choanoeca perplexa P006495
Salpingoeca infusionum P000143
Hartaetosiga gracilis P009096
Hartaetosiga balthica P009124
Microstomoeca roanoka P000728
Salpingoeca rosetta XP_004988131
Barroeca monosierra P008129
B
scarlet
white
MbABCG1
ABCG5, ABCG8
ABCG2
Animals
Choanoflagellates
Holozoans
Fungi
0.5 substitutions/site
