## Supporting Tables for "Choanoflagellates discriminate among *Pseudomonas aeruginosa* alkylquinolones during pathogen response"

**Table 1. HNQ treatment of *M. brevicollis* cultures does not affect feeding bacteria**

| | Treatment | Rep 1 (CFU/mL) | Rep 2 (CFU/mL) | Avg. CFU/mL |
|---|---|---|---|---|
| Experiment 1 | DMSO (control) | 2.6e7 | 2.7e7 | 2.65e7 |
| | HNQ (2.5 μM) | 2.8e7 | 2.75e7 | 2.7125e7 |
| Experiment 2 | DMSO (control) | 1.8e7 | 1.1e7 | 1.45e7 |
| | HNQ (2.5 μM) | 1.9e7 | 1.1e7 | 1.475e7 |

**Table 2. Quantification of secreted AQs during planktonic growth**

| OD=600 | HNQ (μg/mL) | | HHQ (μg/mL) | | PQS (μg/mL) | |
|---|---|---|---|---|---|---|
| | Rep 1 | Rep 2 | Rep 1 | Rep 2 | Rep 1 | Rep 2 |
| 1.2 | 0.008 | 0.004 | 0.030 | 0.028 | 0.052 | 0.042 |
| 1.8 | 0.006 | 0.010 | 0.040 | 0.041 | 0.066 | 0.063 |
| 2.6 | 0.059 | 0.027 | 0.160 | 0.158 | 0.363 | 0.346 |

*strain mPAO1

**Table 3. Quantification of secreted AQs in *pqs* mutant strains (OD=1.2)**

| Strain | AQ per mL (μg/mL) | | |
|---|---|---|---|
| | HNQ | HHQ | PQS |
| *pqsA–* | n.d. | n.d. | n.d. |
| *pqsD–* | n.d. | n.d. | n.d. |
| *pqsE–* | 0.069 | 0.138 | 0.212 |
| *pqsB–* | n.d. | n.d. | n.d. |
| *pqsH–* | 0.050 | 0.071 | 0.027 |

**Table 4. Quantification of AQs in isolated OMVs and supernatant**

| OD=600 | | AQ per mL (μg/mL) | | |
|---|---|---|---|---|
| | | HNQ | HHQ | PQS |
| 3.6 | OMVs | 0.133 | 0.045 | 0.045 |
| | Supernatant | 0.009 | 0.035 | 0.473 |
