## Supplementary material for "Choanoflagellates discriminate among *Pseudomonas aeruginosa* alkylquinolones during pathogen response": Physical data and NMR spectra

Cyclo(Tyr-Pro): white solid powder. HR-ESI-MS *m/z* 261.1232 $[M+H]^+$ calcd. for $C_{14}H_{17}N_2O_3$ at 261.1239 $\Delta_{ppm}$ 2.68. $^1$H NMR (500 MHz, MeOH-$d_4$) $\delta$ 7.04 (d, $J$ = 8.4 Hz, 2H), 6.70 (d, $J$ = 8.4 Hz, 2H), 4.36 (s, 1H), 4.05 (dd, $J$ = 10.8, 6.3 Hz, 1H), 3.55 (dt, $J$ = 11.8, 8.3 Hz, 1H), 3.35 (dt, $J$ = 12.5, 6.6 Hz, 1H), 3.06 (qd, $J$ = 14.2, 4.9 Hz, 2H), 2.10 (dq, $J$ = 11.7, 5.7, 5.2 Hz, 1H), 1.80 (s, 2H), 1.23 (dt, $J$ = 21.4, 10.0 Hz, 1H). $^{13}$C NMR (125 MHz, MeOH-$d_4$) $\delta$ 170.77, 166.96, 157.69, 132.12, 127.62, 116.18, 60.06, 57.91, 45.91, 37.67, 29.40, 22.72.

Cyclo(Phe-Pro): white solid powder. HR-ESI-MS *m/z* 245.1282 $[M+H]^+$ calcd. for $C_{14}H_{17}N_2O_2$ at 245.1290 $\Delta_{ppm}$ 3.26. $^1$H NMR (500 MHz, $CDCl_3$) $\delta$ 7.35 (t, $J$ = 7.3 Hz, 2H), 7.29 (d, $J$ = 7.1 Hz, 1H), 7.23 (d, $J$ = 7.5 Hz, 2H), 5.69 (s, 1H), 4.28 (dd, $J$ = 10.4, 3.0 Hz, 1H), 4.08 (t, $J$ = 7.8 Hz, 1H), 3.69 – 3.52 (m, 3H), 2.79 (dd, $J$ = 14.5, 10.6 Hz, 1H), 2.39 – 2.29 (m, 1H), 2.00 (d, $J$ = 8.0 Hz, 2H), 1.91 (dt, $J$ = 15.3, 8.0 Hz, 1H). $^{13}$C NMR (125 MHz, $CDCl_3$) $\delta$ 169.56, 165.21, 136.06, 129.42, 129.26, 127.70, 59.28, 56.33, 45.60, 36.92, 28.49, 22.69.

2-heptyl-4-(1H)-quinolone (HHQ): white solid powder. HR-ESI-MS *m/z* 244.1691 $[M+H]^+$ calcd. for $C_{16}H_{22}NO$ at 244.1701 $\Delta_{ppm}$ 4.09. $^1$H NMR (500 MHz, MeOH-$d_4$) $\delta$ 8.55 (s, 1H), 8.21 (d, $J$ = 8.2 Hz, 1H), 7.70 (t, $J$ = 7.7 Hz, 1H), 7.59 (d, $J$ = 8.4 Hz, 1H), 7.40 (t, $J$ = 7.6 Hz, 1H), 6.24 (s, 1H), 2.77 – 2.68 (m, 2H), 1.78 (p, $J$ = 7.6 Hz, 2H), 1.40 (m, 3H), 1.36 – 1.27 (m, 5H), 0.90 (t, $J$ = 6.8 Hz, 3H). $^{13}$C NMR (125 MHz, MeOH-$d_4$) $\delta$ 180.67, 170.23, 157.17, 141.62, 133.41, 125.98, 125.49, 125.07, 119.07, 108.83, 35.00, 32.87, 30.19, 30.10, 23.66, 14.38.

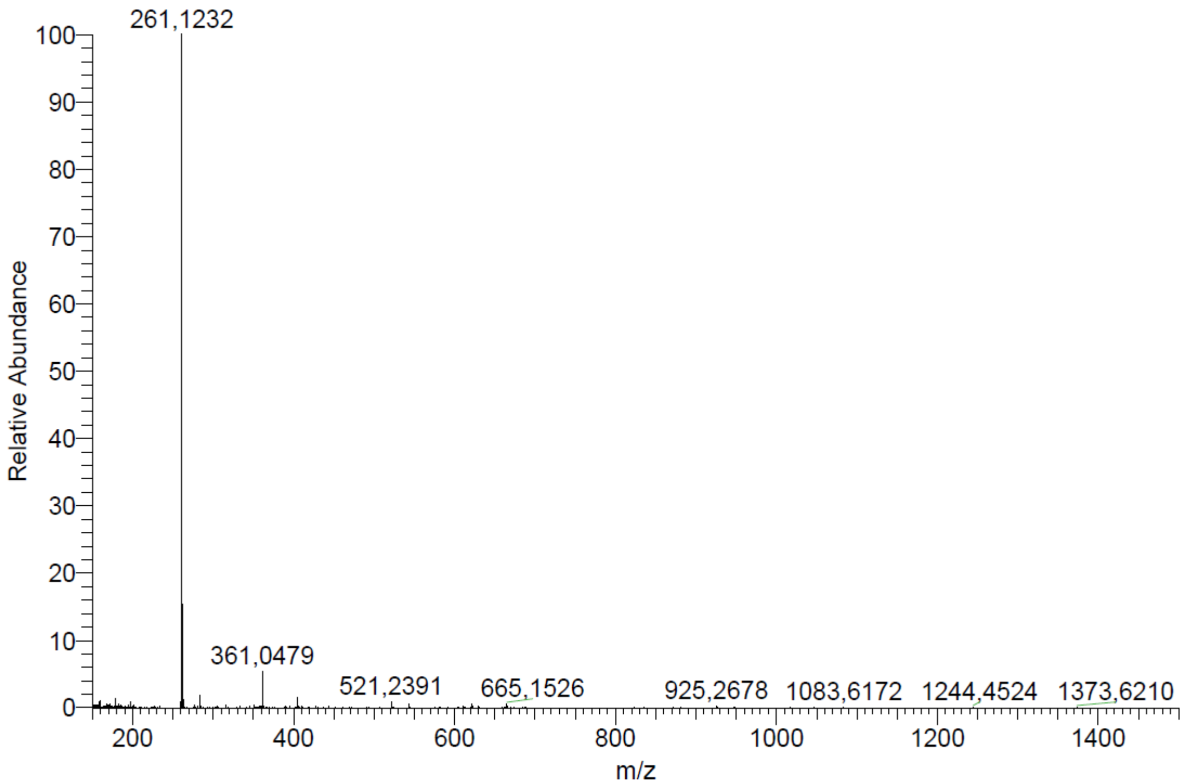


ESI-HRMS spectrum of Cyclo(Tyr-Pro)

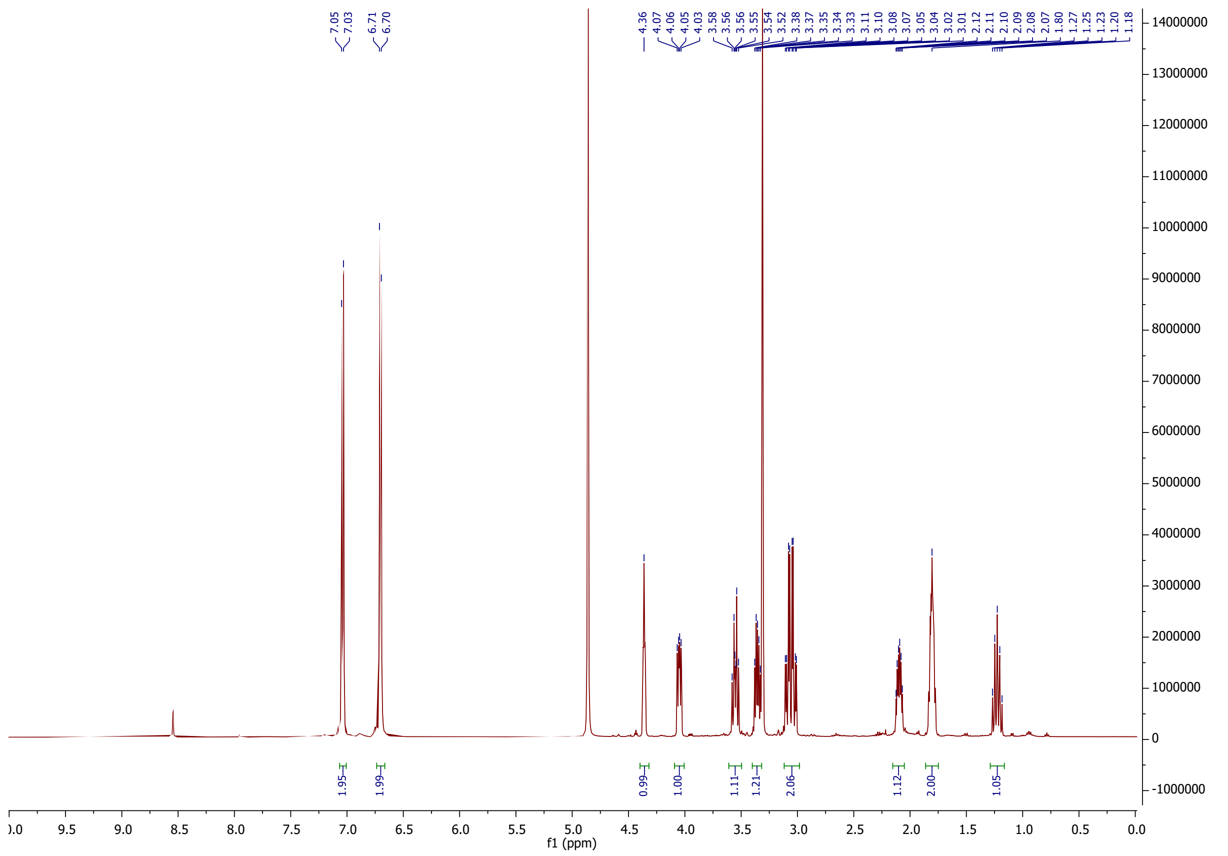

$^1H$ NMR spectrum of Cyclo(Tyr-Pro) (500 MHz, MeOH-$d_4$)

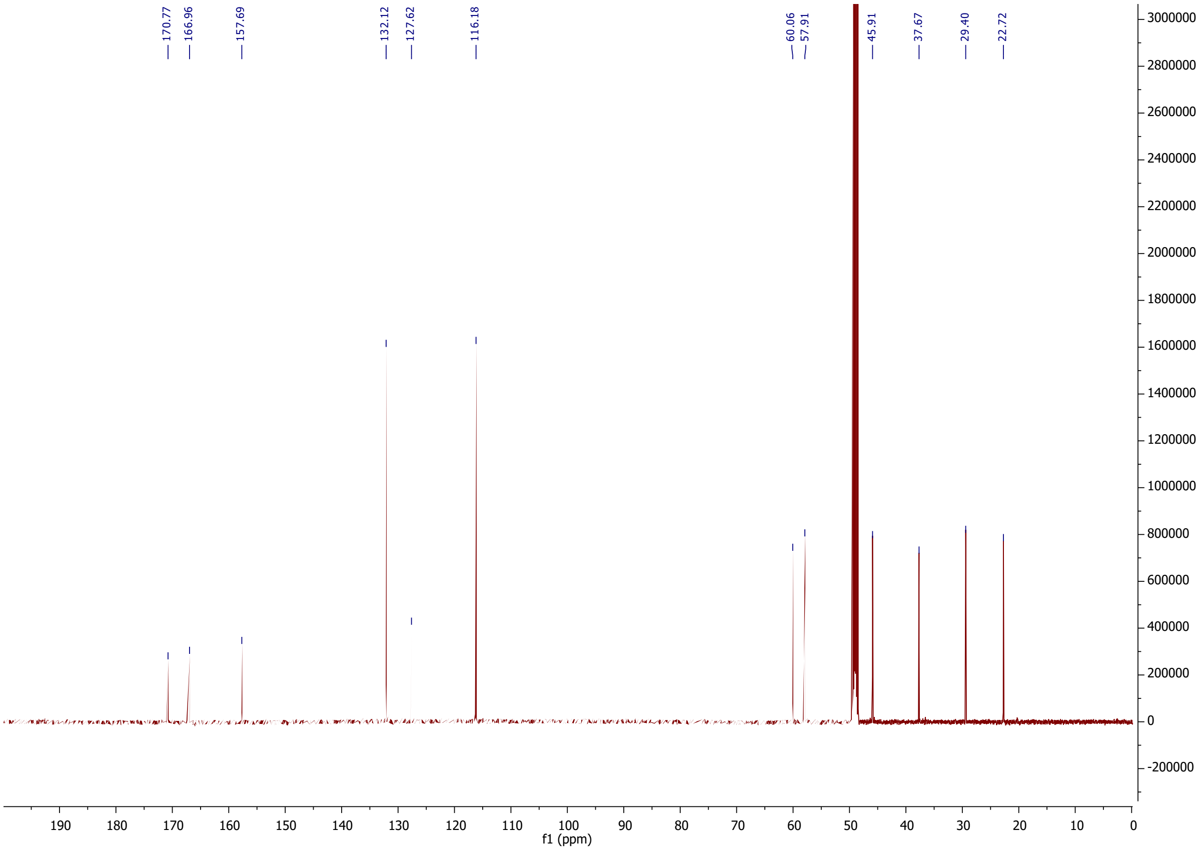


$^{13}C$ NMR spectrum of Cyclo(Tyr-Pro) (125 MHz, MeOH-$d_4$)

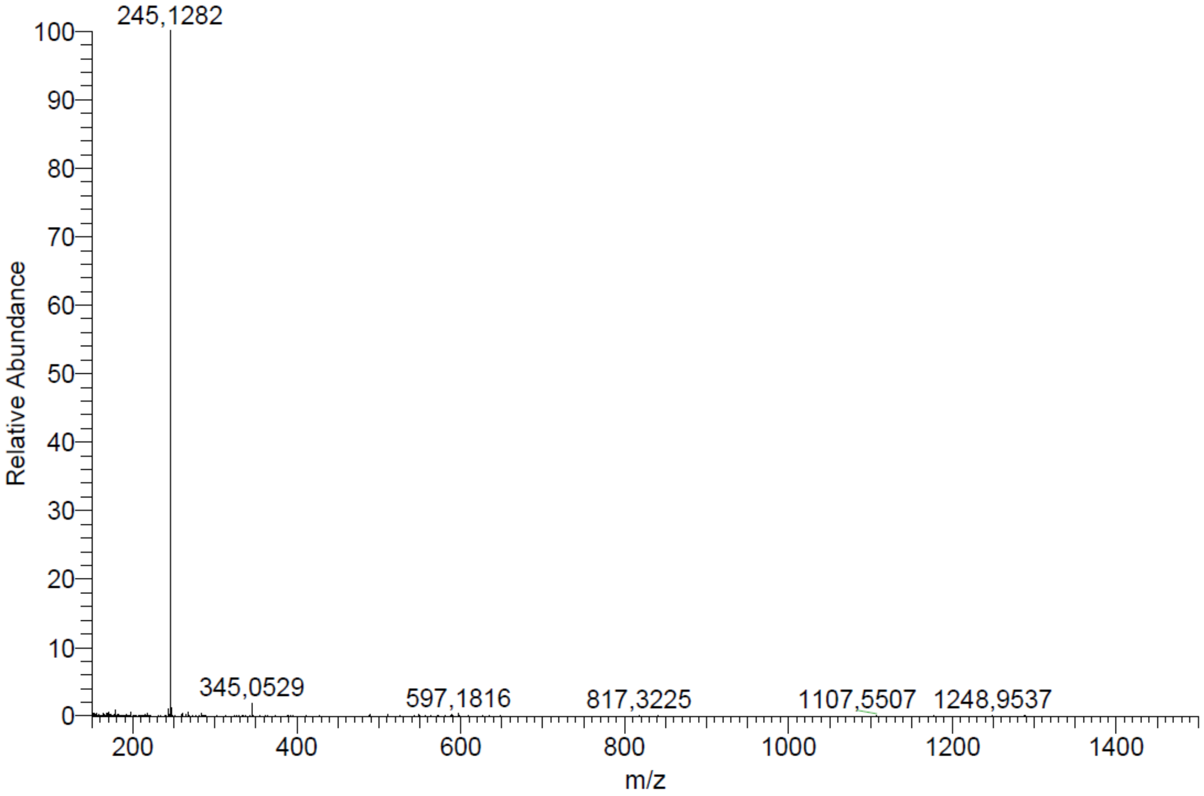


ESI-HRMS spectrum of Cyclo(Phe-Pro)

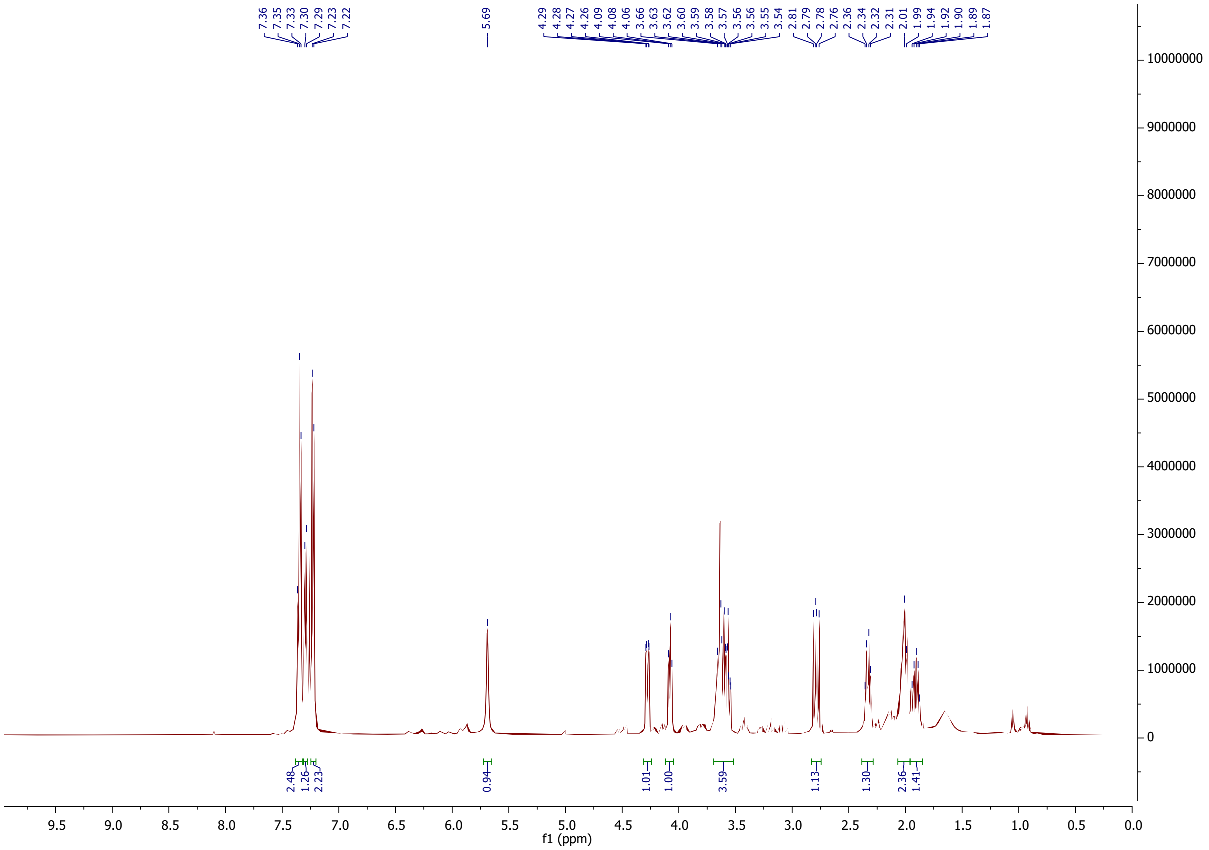


$^1H$ NMR spectrum of Cyclo(Phe-Pro) (500 MHz, $CDCl_3$)

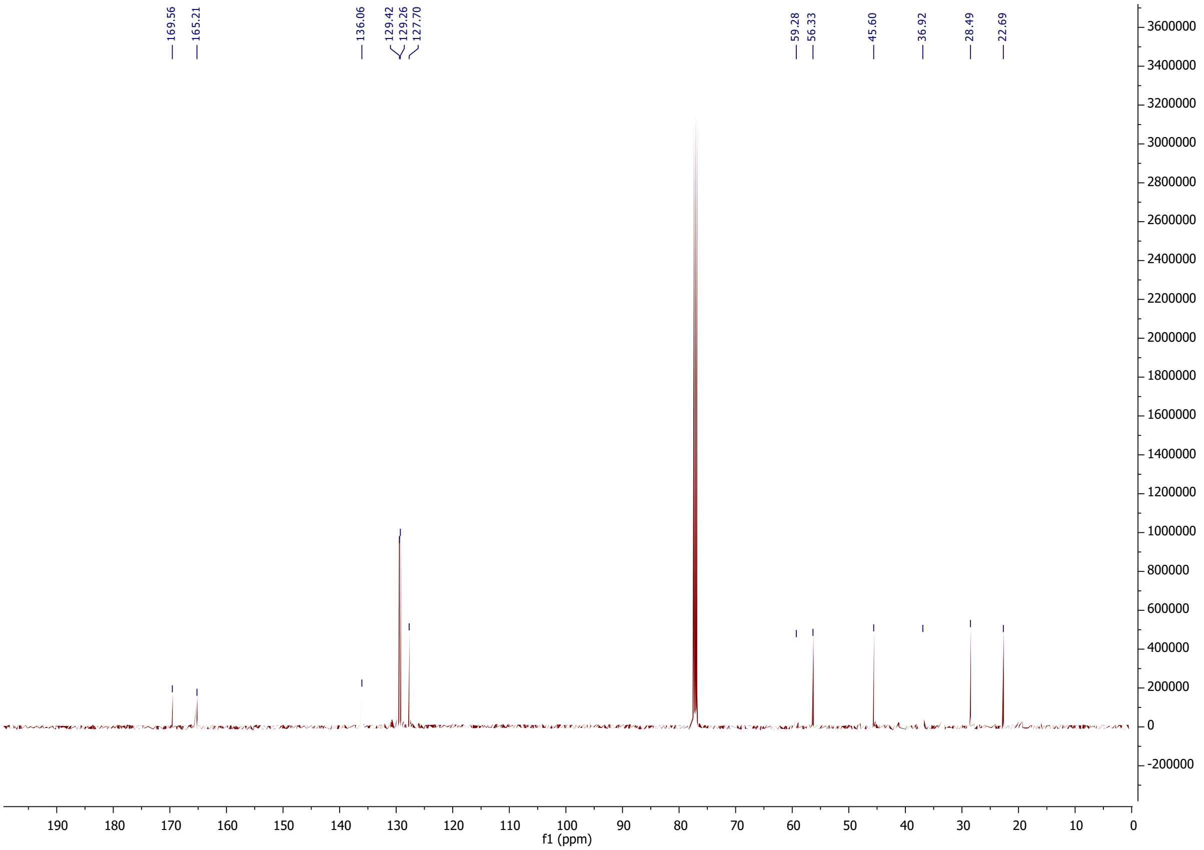


$^{13}C$ NMR spectrum of Cyclo(Phe-Pro) (125 MHz, $CDCl_3$)

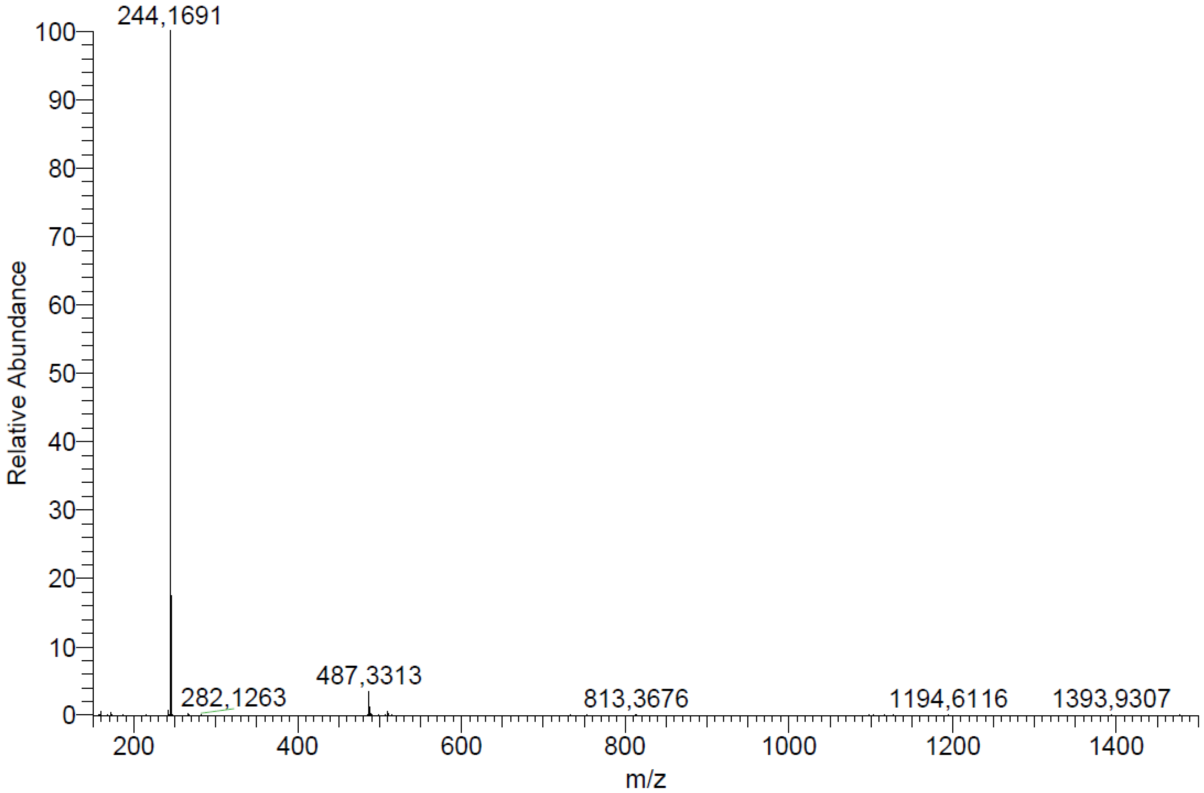


ESI-HRMS spectrum of HHQ

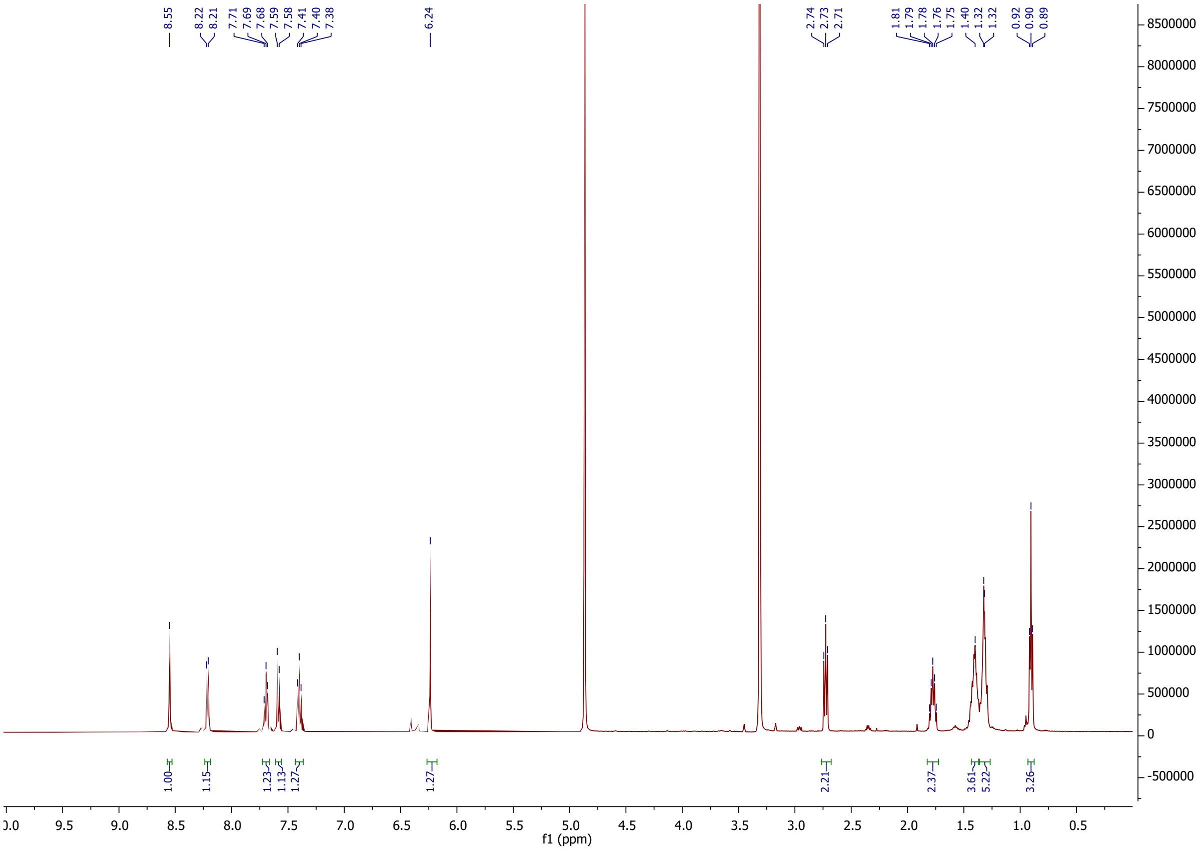


$^{1}$H NMR spectrum of HHQ (500 MHz, MeOH-$d_4$)

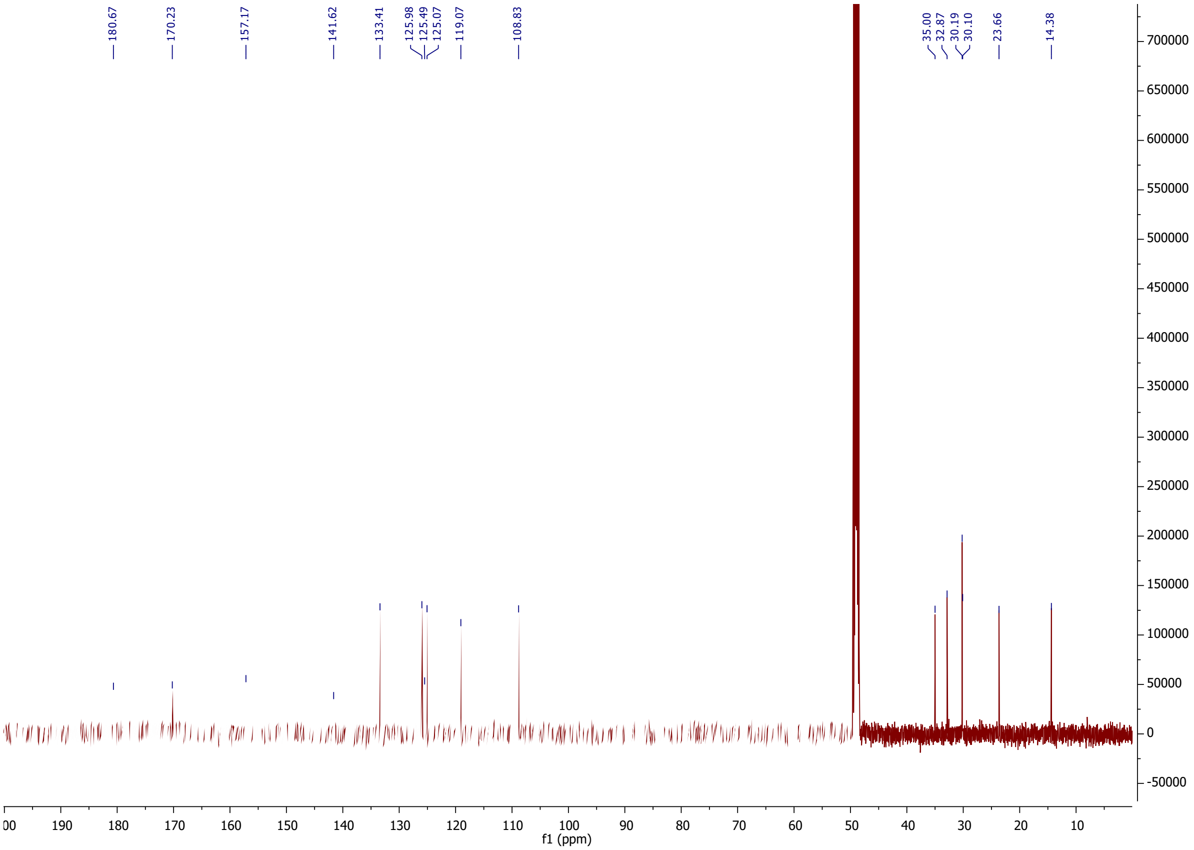


$^{13}$C NMR spectrum of HHQ (125 MHz, MeOH-$d_4$)
