## Supplementary material for "Choanoflagellates discriminate among *Pseudomonas aeruginosa* alkylquinolones during pathogen response": Methods

#### Cell Culture

##### Choanoflagellate cell culture

*M. brevicollis (*American Type Culture Collection [ATCC], Manassas, VA; Cat. No. PRA-258) was co-cultured with feeding bacterium *P. shioyasakiensis* in a seawater based media enriched with glycerol, yeast extract, peptone and cereal grass (10% CGM3, 4% SWC). *S. rosetta* co-cultured with *E. pacifica* feeding bacteria[1] (American Type Culture Collection [ATCC], Manassas, VA; Cat. No. PRA-390) was cultured in a seawater based media enriched with glycerol, yeast extract, peptone and cereal grass (10% CGM3, 2.5% SWC). *S. rosetta* rosette formation was induced by adding OMVs isolated from *A. machipongonensis* bacteria[2]. *C. flexa* (environmental isolate, species described in[3]) was cultured alongside a diverse assortment of co-isolated environmental bacteria in 25% CGM3. *M. fluctuans* (American Type Culture Collection [ATCC], Manassas, VA; Cat. No. 50635) was cultured in 10% FCGM3 2.5% FWC. All choanoflagellate species were passaged in vented cultured flasks and incubated at 22°C and 60% relative humidity or at 16°C.

##### Mammalian cell culture

THP-1 (American Type Culture Collection [ATCC], Manassas, VA; Cat. No. TIB-202) and A549 (American Type Culture Collection [ATCC], Manassas, VA; Cat. No.CRM-CCL-185) cell lines were maintained at 37 °C, 5% $CO_2$ and subcultured according to ATCC recommended procedures. THP-1 cells were cultured in RPMI-1640 containing L-glutamine and sodium bicarbonate (Sigma Aldrich), supplemented with 10% (v/v) heat-inactivated fetal bovine serum (FBS; Sigma Aldrich), 1% (v/v) penicillin–streptomycin, 1% (v/v) sodium pyruvate (GIBCO). A549 cells were cultured in high-glucose DMEM (GIBCO) with 10% (v/v) FBS.

#### HPLC fractionation of *P. aeruginosa* conditioned media and bioassays

Single *P. aeruginosa* (mPAO1 or PW6886, obtained from the PAO1 Two-Allele Transposon Mutant Library) colonies were inoculated into LB and grown at 30 °C with shaking at 225 rpm. Overnight cultures were sub-cultured into 500mL fresh medium ($OD_{600}$=0.05-0.1) and grown for 16 hours. Bacterial cells were pelleted by centrifugation in polypropylene bottles at 5,000 x g for 30 min (up to 9,200 x g for ≥75% filled vessels). Supernatant was decanted from the bacterial cell pellet and filtered twice through 0.22 μm sterilizing filters to yield conditioned media.

Conditioned media (750mL) was extracted with 1:1 EtOAc three times in a separative funnel. Organic phases were pooled and dried under vacuum to give 104 mg of crude extract. Crude extract was subjected to semi-preparative reverse-phase HPLC (Phenomenex Luna C18(2) 250 x 10 mm 5 μm, 100 Å column,) using the following gradient: 0–5 min, 10% B; 5–36 min, 10%–100% B; 36–44 min, 100% B (A: $H_2O$ + 0.1% Formic acid; B: $CH_3CN$ + 0.1% Formic acid) with a flow rate of 3 mL/min to obtain 16 fractions, three of which yielded pure compounds: Cyclo(Tyr-Pro) (6.2 mg), Cyclo(Phe-Pro) (3.9 mg) and 2-heptyl-4(1H)-quinolone (HHQ) (0.6 mg). Each HPLC fraction was resuspended to 1 mg/mL in DMSO prior to performing 1:5 serial dilutions. To assess bioactivity, 5 µL of each diluted fraction (beginning at 1:5) was added to 145 µL *M. brevicollis* (cell density of 1x$10^5$ cells/mL) in a 96-well plate format. Plates were incubated at 16 °C for 20 hours, and cell counts were determined using a hemocytometer.

**Isolation and extraction of *Pseudomonas* conditioned media and bioassays**

*P. aeruginosa* alkylquinolone biosynthesis mutants: *P. aeruginosa* parental strain and transposon mutants we obtained from the PAO1 Two-Allele Transposon Mutant Library[4] and confirmed by PCR and Sanger sequencing. To generate conditioned media, single colonies were inoculated into LB and grown at 37 °C with shaking at 225 rpm. Overnight cultures were sub-cultured into 50mL fresh medium ($OD_{600}$=0.05-0.1) and grown until $OD_{600}$=1.2.

Diverse *Pseudomonas* species: To generate conditioned media, single isolated colonies were picked and inoculated into LB ang grown at 30 °C with shaking at 250 rpm. Overnight cultures were subcultured into 250 ml fresh LB ($OD_{600}$=0.05) and grown until $OD_{600}$=1.2. Bacterial cells were pelleted by centrifugation at 3,290 x g for 30 min (for smaller volumes) or at 10,000 x g for 20 min. Supernatant was decanted from the bacterial cell pellet and filtered twice through 0.22 µm sterilizing filters to yield conditioned media. Conditioned media was then used in bioassays or for further metabolic profiling.

Extraction procedure: All conditioned media was extracted with EtOAc. The organic extract was dried under vacuum, dissolved in MeOH with a final concentration of 50 µg/mL, and analyzed by UHPLC-HRMS.

Bioassays: Choanoflagellates cell densities were determined using a hemocytometer, then cells were diluted to 1x$10^5$ cells/ml and subsequently treated with 5% (v/v) conditioned media in a 24-well plate format. Plates were incubated at 22 °C for 20 hours, and cell counts were determined using a hemocytometer.

**UHPLC-HRMS analysis and NMR**

All media ingredients, reagents and chemicals were purchased as follows: Methanol (Th. Geyer, Renningen); water for analytical and preparative HPLC (VWR International GmbH, Dresden); formic acid (VWR International GmbH, Dresden); acetonitrile (Th. Geyer, Renningen); standard and $CDCl_3$ and MeOH-$d_4$ (Sigma Aldrich, USA).

UHPLC-HRMS measurements were carried out on a Vanquish Flex UHPLC system (Thermo Fisher Scientific) combined with an Orbitrap Exploris 120 mass spectrometer equipped with a HESI source (Thermo Fisher Scientific). Metabolites were separated using reverse phase liquid chromatography at 40 °C using a Kinetex® C18 column (50 × 2.1 mm, particle size 1.7 µm, 100 Å, Phenomenex) preceded by a C18 SecurityGuardTM ULTRA guard cartridge (2.1 mm, Phenomenex). Mobile phases consisted of $H_2O$ + 0.1 % formic acid (A) and acetonitrile (ACN) + 0.1 % formic acid (B). 5 µL sample, concentrated at 50 µg/mL, was injected into a gradient as follows: 0-1 min, 5 % B; 1-10 min, 5-97 % B; 10-12 min, 97 % B; 12-13 min, 97-5 % B; 13-15 min, 5 % B at a constant flow rate of 0.3 mL/min. Data dependent acquisition of $MS^2$ spectra was performed in positive mode. HESI parameters were set to 50 AU sheath gas flow, 13 AU auxiliary gas flow, 1 AU sweep gas flow, 3.4 kV (+) spray voltage, 300 °C vaporizer temperature and 320 °C ion transfer tube temperature. $MS^1$ full scan parameters were set to data type-centroid, *m/z* 150-1,500 scan range, resolving power 60,000 at *m/z* 200, 1 micro-scan, 100 ms max. injection time, 1E6 automated gain control, 70 % RF lens, dynamic exclusion filter-auto and isotope exclusion filter-assigned. Up to 4 $MS^2$ spectra per $MS^1$ survey scan were recorded with the following parameters, data type-centroid, scan range-auto, resolving power 30,000 at *m/z* 200, 1 micro-scan, 100 ms max. injection time, 1E5 automated gain control, 1.2 *m/z* isolation window, collision energy type- normalized with a stepwise increase from 20 to 30 to 40 %.

**Calibration curve and quantification by UHPLC-HRMS**

2-Nonylquinolin-4-one, 4-hydroxy-2-heptylquinoline (HHQ), and 2-heptyl-3-hydroxy-4-quinolone (PQS) were identified by comparison with commercial standards using the UHPLC-HRMS method described above. Quantification was performed using a calibration curve prepared with 2-nonylquinolin-4-one at concentrations of 0.1, 0.5, 1, 5, and 10 µg/mL in methanol ($R^2$ = 0.9992). All three compounds were quantified using this calibration curve based on structural similarity.

**Feature-based molecular networking**

The raw MS2 data files were converted to mzML using MSconvert[5,6] . The MZmine[7] version 2.53 was then used for pre-processing. A centroid mass detection of MS1 at a noise level of 1.0E6 and MS2 noise level of 1.0E3, generated a mass list. The m/z tolerance was set to 0.01 Da. The feature list was used to build chromatograms using ADAP. The minimum group number of scans was set at 5, the group intensity threshold was set at 1.0E6 and the minimum highest intensity was set at 3.0E6. Deconvolution of the feature lists was done with the Baseline cut-off algorithm. The feature lists were deisotoped and the features were aligned. Finally, duplicates were filtered, whereas fragments and adducts were searched. The resulting aligned list of features was exported in a mgf file along with their feature quantification table in csv format. These data as well as the mzML files were uploaded onto the feature-based molecular network (FBMN) page of GNPS2[8,9] Networks were generated using six minimum matched ion fragments, a minimum cluster size of two and a cosine score of 0.7. The remaining parameters were kept at default. The molecular network was visualized on Cytoscape version 3.9.1.[10] All data set used for molecular networks have been submitted to MassIVE repository.

**Statistics**

Principal Component Analysis (PCA) and Hierarchical Clustering Dendrogram were performed by using MetaboAnalyst[11]. Before statistics analyses, data was interquartile data filtered and pareto scaled by using variance transformation.

**Commented [MD1]:** Pang, Z. *et al.* MetaboAnalyst 6.0: towards a unified platform for metabolomics data processing, analysis and interpretation. *Nucleic Acids Res* **52**, W398–W406 (2024).

The NMR spectra of the compounds were collected using an NMR spectrometer outfitted with a Bruker CryoPlatform ($CDCl_3$, MeOH-$d_4$, 500 MHz). MestReNova ver. 12.0.1 was used to process the data for the NMR spectra.

***P. aeruginosa* Outer Membrane Vesicles (OMVs)**

OMV Isolation: Single *P. aeruginosa* colonies were inoculated into LB and grown at 30 °C with shaking at 225 rpm. Overnight cultures were sub-cultured into fresh medium ($OD_{600}$=0.05-0.1) and grown for 16 hours. Bacterial cells were pelleted by centrifugation in polypropylene bottles at 5,000 rpm for 30 min (up to 10,000 rpm for ≥75% filled vessels). Supernatant was decanted from the bacterial cell pellet and filtered twice through 0.22 um sterilizing filters to yield conditioned media. OMVs were isolated from conditioned media by centrifugation at 36,000

RPM (using a Beckman Type 45Ti fixed angle rotor) for 3 hours at 4 °C. Supernatant was decanted. Pellets were briefly dried at RT prior to being resuspended in 50 mM HEPES (pH7.4) for use in downstream bioassays and UHPLC-HRMS analysis.

Bioassays: Choanoflagellates cell densities were determined using a hemocytometer, then cells were diluted to $1x10^5$ cells/ml in a 24-well plate format. Resuspended OMVs were diluted such that 1 µL OMVs was equivalent to 1 mL conditioned media. Cells were then incubated with 5% v/v *P. aeruginosa* conditioned media, 5% v/v Supernatant, or 0.005% v/v OMVs. Plates were incubated at 22 °C for 20 hours, and cell counts were determined using a hemocytometer.

Imaging: For imaging experiments, OMVs were stained with a lipophilic dye (FM 4-64FX or FM 1-43FX, Invitrogen) following the manufacturer's recommendations. Excess dye was removed using a Zeba desalting column and OMVs were resuspended to original volume. Choanoflagellates cell densities were determined using a hemocytometer, then cells were diluted to $1x10^5$ cells/ml. 0.005% v/v OMVs were added to cells for 30 minutes, prior to fluorescent and DIC imaging.

Widefield images were acquired on a Nikon TI2-E inverted microscope with DIC, multiline LED fluorescence, an ORCA-Fusion BT camera, and a 60x Water objective (NA – 1.2; WD - 0.18 mm). Super resolution images were acquired on a Nikon AXR-NSPARC system equipped with a Plan Apochromat λ D 60x oil objective (NA – 1.2; WD - 0.18 mm). Fluorescent images were acquired using a FN25 galvano scanner, 488 nm and 568 nm lasers, and a 0.16 µM step size. At least 20 cells were imaged per condition. Following acquisition, Z-stacks were 3D aligned and deconvolved using Nikon Elements software. Imaris software was used to visualize maximum intensity projections and 3D object rendering with transparency, allowing us to visualize intracellular bacteria and bacteria at the collar membrane.

**Experiments using synthesized or isolated AQs**

All quinolones were resuspended to 2mM. Serial dilutions of all synthesized quinolones were performed in DMSO.

Choanoflagellates: Choanoflagellates cell densities were determined using a hemocytometer, then cells were diluted to either $1x10^4$ cells/mL (*M. fluctuans)* or $1x10^5$ cells/mL (*M. brevicollis, S. rosetta, C. flexa*) in growth media. Cells were plated into 96 well plates, with 147 µL cells/well. Each well was then treated with 3 µL of the quinolone compounds or DMSO. Cell counts were obtained after 22 hours (for *M. brevicollis* and *C. flexa*), 42 hours (for *S. rosetta*), or 67 hours (for *M. fluctuans)*.

*M. brevicollis* HNQ Viability Time Course:

*M. brevicollis* was grown for 24 hours, and then diluted to $5x10^5$ cells/mL in growth media (10%CGM3, 4%SWC) in 24-well plates. Cells were treated with indicated concentrations of HNQ or DMSO, and counted after 2, 7, 16, and 24 hours. Movies were acquired using a 60x water immersion objective on a Nikon TI2-E inverted microscope with DIC and ORCA-Fusion BT camera at 1, 3, and 5 hours post-treatment.

A549 cells: A549 Cells were seeded at $1x10^5$ cells/mL in a 96-well plate ($1x10^4$ cells/well), and allowed to adhere for 24 h prior to treatment. Immediately prior to treatment, quinolone compounds were further diluted 1:50 in complete growth medium, and 10 µL was added to each well. After 24 hours, growth medium was removed, and cells were washed with 1X PBS and stained with Hoechst 33342 (1:2000 in 1X PBS; Invitrogen, Carlsbad, CA). Cell number was quantified using automated cell imaging on a Nikon TI2-E inverted microscope with DIC, multiline LED fluorescence, an ORCA-Fusion BT camera. Each well was imaged using 4x4 tiling, and clarified using Nikon Elements software. Binary images were then generated using Fiji[12] software. Finally, automated cell counts were generated via the Analyze Particles function in Fiji.

THP-1 cells: THP-1 cells were seeded at $2x10^5$ cells/mL in a 96-well plate ($2x10^4$ cells/well) and allowed to grow for an additional 24 hours. Immediately before treatment, quinolone compounds were further diluted 1:10 in MeOH, and 2.5 µL was added to each well. Cell densities were quantified 24 hours after treatment, using either a hemocytometer or LUNA cell counter.

**Phagocytosis Imaging**

Single colonies of GFP labeled mPAO1 or BW25113 were inoculated into LB and grown shaking at 37 C. When OD600=0.4-0.5, bacteria were diluted to a concentration of $10^7$ cells/mL in water.

*M. brevicollis* stably expressing LifeAct-mCherry was grown for 24 hours, washed twice by centrifugation to reduce feeding bacteria, and resuspended to a final concentration of $10^6$ cells/mL in artificial sea water. 200 µl aliquots were transferred into microcentrifuge tubes and inoculated with bacteria at MOI=10. Choanoflagellates were allowed to feed for 2.5 hours at room temperature, then transferred to an 18-well chamber slide (Ibidi) for live imaging.

Cells were imaged on a Nikon AXR-NSPARC system equipped with a Plan Apochromat λ D 60x oil objective (NA – 1.2; WD - 0.18 mm). Fluorescent images were acquired using a FN25 galvano scanner, 488 nm and 568 nm lasers, and a 0.16 µM step size. At least 20 cells were imaged per condition. Following acquisition, Z-stacks were 3D aligned and deconvolved using Nikon Elements software. Imaris software was used to visualize maximum intensity projections and 3D object rendering with transparency, allowing us to visualize intracellular bacteria and bacteria at the collar membrane.

**RNAseq**

*M. brevicollis* was grown for 24 hours in 3-layer flasks (Corning Life Sciences, Tewksbury, MA) at 16 °C. Cells were washed three times by centrifugation (2,000 x *g*) in artificial seawater (ASW), and then resuspended to a concentration of $1x10^7$ cells/mL in growth media (10% CGM3, 4% SWC).

Two technical replicates, each consisting of $5x10^7$ cells, were pooled for each biological replicate. Samples were either left untreated, incubated with 1.25 µM HNQ, or incubated with 5% (v/v) PAO1 conditioned medium (collected at $OD_{600}$=2.4). Samples were incubated at room temperature with gentle nutation for 3 hours, prior to pelleting by centrifugation at 3,290 x g for 15 min at 4 °C.

Total RNA was extracted using the RNAqueous kit (Thermo Fisher Scientific), following the manufacturer's instructions, with the following modifications: buffer volumes were doubled, and RNA was eluted in TE buffer. mRNA libraries were prepared using Illumina library prep kits and a starting concentration of 2 µg RNA, and sequenced on an Illumina NovaSeq platform (2x150bp sequencing reads, 25M Reads) (SeqCenter). Quality control and adapter trimming was performed with bcl-convert, and read mapping was performed via STAR[13] to the *M. brevicollis* genome[14]. Feature quantification was performed using RSEM[15]. Read counts were normalized using edgeR's Trimmed Mean of M values (TMM) algorithm[16] . Subsequent values were then converted to counts per million (CPM).

**Shotgun proteomics**

Cell growth and sample preparation: *M. brevicollis* was grown for 24 hours in 3-layer flasks (Corning Life Sciences, Tewksbury, MA) at 16 °C. Cells were washed three times by centrifugation (2,000 x *g*) in artificial seawater (ASW), and then resuspended to a concentration of $1x10^7$ cells/mL in growth media (10% CGM3, 4% SWC). $1x10^8$ cells/mL were used for each of the three biological replicates per condition. To assess changes in protein abundance following HNQ treatment, cells were treated either with 1.25 µM HNQ or DMSO (control) for 5 hours. Cells were then pelleted at 2,300 x *g* for 5 min, and supernatant was removed. Pellets were resuspended in 3 mL ice cold HEPES Lysis Buffer (50 mM HEPES pH 7.4, 150 mM NaCl, 50 mM MgSO4, 1 mM EGTA, 1% NP-40, 0.1% SDS, 1mM DTT, .1 mM PMSF, EDTA-free protease inhibitor, PhosSTOP) and incubated on ice for 10 mins with frequent mixing. Benzonase (25U/µL) was then added to the lysate, which was incubated on ice for an additional 30 minutes. To clarify the lysate, samples were centrifuged at 20,000 x g for 20 minutes at 4°C. Clarified supernatant was transferred into low-bind centrifuge tubes. ~50µL was aliquoted for DC Bradford Assay, and the remaining lysate was stored at -80°C such that all biological replicates could be processed simultaneously for proteomics.

Lysates were prepared for mass spectrometry with the solution-based combinative sample preparation (CSP) method of Lin et al[17]. Briefly, SDS was added to 200 µL of lysate (2 µg/µL protein) to a final concentration of 2%, and samples were heated at 95 °C for 5 min. Proteins were precipitated overnight at 4 °C with six volumes of acetone pre-chilled to −20 °C, pelleted at 16,000 × g for 15 min at 4 °C, and washed twice with pre-chilled acetone. Dried pellets were resuspended in 100 µL of 1% sodium deoxycholate (SDC) in 50 mM ammonium bicarbonate by bath sonication for 20 min. Proteins were reduced with TCEP at 56 °C for 45 min, alkylated with 25 mM iodoacetamide for 45 min at room temperature in the dark, and quenched with 12 mM DTT. Proteins were then digested overnight at 37 °C with 1.6 µg of MS-grade trypsin (Pierce #90058). Digestion was stopped and SDC was precipitated by adding formic acid to a final concentration of 1%, and the precipitate was removed by centrifugation at 16,000 × g for 10 min at 4 °C. The supernatant was passed through a pre-washed Vivaspin 500 10 kDa MWCO filter (Sartorius) and dried by vacuum centrifugation. Peptides were resuspended in 20 µL of 0.1% trifluoroacetic acid, desalted with a ZipTip (Millipore #ZTC18S096), dried again, and resuspended in 10 µL of 5% acetonitrile (ACN), 0.1% formic acid. Each sample was injected twice as technical replicates, for a total of 24 LC-MS/MS runs.

**LC-MS/MS and protein identification**

Peptides were separated on a Dionex UltiMate 3000 RSLCnano UHPLC system (Thermo Scientific) fitted with a C18 trap to an EASY-Spray PepMap RSLC C18 analytical column (Thermo Scientific), using a 60-min gradient of 3–45% ACN. Eluted peptides were ionized by nano-electrospray and analyzed on an Orbitrap Fusion Lumos Tribrid mass spectrometer

(Thermo Scientific). Data were acquired with a 75-min data-dependent method in top-speed mode, in which precursor ions were fragmented by high-energy collisional dissociation (HCD; stepped collision energies of 27, 30, and 33%) and fragment ions were detected in the ion trap.

Raw files were processed in Proteome Discoverer 2.5 (Thermo Scientific). Each file was first searched separately with a processing workflow based on the PWF_Tribrid_Basic_SequestHT template (spectrum recalibration, Sequest HT, and Percolator). The search database comprised the *M. brevicollis* UniProt reference proteome (UP000001357; 9,188 entries, downloaded January 16, 2024), 545 hypothetical proteins from (bacterial species/strain and genome source), and 379 common contaminant proteins[18]. Search parameters were full trypsin specificity with up to two missed cleavages, precursor and fragment mass tolerances of 10 ppm and 0.6 Da, respectively, carbamidomethylation of cysteine as a static modification, and methionine oxidation and protein N-terminal acetylation, Met-loss, or Met-loss + acetylation as dynamic modifications. The two injections of each biological replicate were then combined in a consensus workflow (merge mode "Globally by Search Engine Type"), yielding one result per biological replicate. Only high-confidence peptide-spectrum matches (PSMs) and peptides at 1% false discovery rate (FDR) were retained. Proteins were assigned high (1% FDR) or medium (5% FDR) confidence, and protein groups were formed under the strict parsimony principle.

**Mass spectrometry data analysis**

Protein abundance was compared across conditions by spectral counting. For each biological replicate, the spectral count of a protein was defined as the number of PSMs assigned to it as master protein in the consensus result. Counts from the biological replicates were combined into one table, and contaminant and bacterial proteins were removed. Proteins with at least two PSMs in at least two samples were retained, and differential abundance of each treatment relative to ASW was tested with the RUVr edgeR quasi-likelihood model in Degust[19], using one factor of unwanted variation (k = 1) and trimmed mean of M-values (TMM) normalization.

**MbABCG1 (*Monbrdraft_34196*) Protein Domain Identification and Phylogenetic Analysis**

We first analyzed the protein sequence of MONBRDRAFT_34196 for conserved protein domains using the HMMER[20] web server. Next, to identify MONBRDRAFT_34196 orthologs and compare it to characterized ABCG transporters, we ran BLASTP (v2.13.0+) against all opisthokont proteomes and transcriptome-derived proteins available in EukProt V3[21]. This returned 507 hits from 114 taxa. Next, near-identical sequences from the same taxon (more than 80% of 8-mers shared) were collapsed to one representative to remove isoforms. The final data set contained 335 sequences from approximately 100 taxa. Sequences were aligned with MAFFT v7.505[22], and maximum-likelihood trees were inferred with IQ-TREE[23] v2.0.7 (LG+F+G4 without model selection, 1000 ultrafast bootstrap replicates. Alternative placements of MbABCG1 were tested by constrained ML searches in IQ-TREE on a 154-sequence subset of the alignment.

***M. brevicollis* genetics and transgenics**

***M. brevicollis* CRISPR-Cas9 Genome Editing**

To disrupt the *MbABCG1* locus, we inserted a terminator sequence in the first exon of *MbABCG1*. First, we designed a 20 bp sgRNA (ordered from IDT) using the Cas9 extended recognition motif (5'-HNNGRSGGH-3') which targeted a cut site 115 bp into Exon 1 of

*MbABCG1*. We designed a dsDNA repair template using the following approach: (1) We generated a reusable "terminator" cloning vector containing a terminator sequence followed by a puromycin resistance cassette driven by the *M. brevicollis* EFL promoter sequence. (2) We identified ~50 bp locus-specific forward and reverse primers to serve as homology arms. The forward primer included *MbABCG1* sequence immediately upstream of the PAM site, while the reverse primer included *MbABCG1* sequence immediately downstream of the PAM site. (3) To generate a repair template, we PCR amplified from the terminator cloning vector using the locus specific forward and reverse primers.

To generate an endogenously tagged C-terminal fusion protein of MbABCG1, we inserted a short linker and fluorescent marker (StayGold) in-frame at the 3' end of the gene coding sequence of *MbABCG1*. We identified a 3'-adjacent Cas9 "NGG" PAM site within the coding region of *MbABCG1*, and designed a 20 bp sgRNA (IDT) immediately upstream of this PAM. We then generated a dsDNA repair template using the following approach: (1) First, we generated a reusable *Staygold* cloning vector containing (i) a flexible linker sequence (GGSGGS), (ii) the *StayGold* sequence codon-optimized for *M. brevicollis,* (iii) a P2A self cleaving peptide, (iv) the puromycin resistance cassette. (2) We then designed ~50 bp locus-specific forward and reverse primers to serve as homology arms. The forward primer included *MbABCG1* coding sequence immediately upstream of the PAM site, while the reverse primer included the endogenous *MbABCG1* stop codon and subsequent 3'UTR sequence. (3) To generate a repair template, we PCR amplified from the *Staygold* cloning vector using the locus specific forward and reverse primers.

Both the terminator and StayGold dsDNA repair templates were amplified using KOD polymerase. PCR amplicons were purified using a Qiagen PCR purification kit and eluted in 25 µL nuclease-free water to yield 8-12 µg dsDNA. Eluant was evaporated on a heat block to reduce the volume of repair template to ~5 µL before transfection. .

CRISPR reactions were transfected and recovered following the protocol from Woznica et al. (2022), with the following changes: (1) sgRNA guides (already complexed with tracrRNA) were ordered from IDT; (2) repair oligos were dsDNA and amplified via PCR.

To select for puromycin-resistant cell lines, cells were expanded into a 3mL volume in 6-well plates and treated with 300 µg/mL puromycin. 5 days post-selection, cells were genotyped at the population level at the locus of interest. If a successful edit was visible by PCR genotyping, cells underwent limiting dilution to isolate clonal populations for further sequencing and downstream analysis.

**Generating Stable *M. brevicollis* Expression Strains**

Plasmids to stably express LifeAct::mCherry and MbABCG1::mTFP in *M. brevicollis* were designed following methods published in Booth et al. (2018)[24] and Woznica et al. (2021)[25].

<u>LifeAct::mCherry</u>: To tune levels of LifeAct::mCherry in the cell, we generated two reporter constructs driven by distinct *M. brevicollis* promoter sequences: the actin (Monbrdraft_37852) promoter and a ribosomal subunit promoter (Monbrdraft_35363). For both constructs, the LifeAct::mCherry sequence is immediately followed by the 3'UTR from *M. brevicollis* actin. The LifeAct::mCherry plasmids also contain the Pac puromycin resistance gene for selection.

MbABCG1::mTFP: We first amplified the CDS of *Monbrdraft_34196* from *M. brevicollis* cDNA. This sequence was then subcloned using Gibson assembly into an *M. brevicollis* expression construct downstream of a 5' actin promoter, and upstream of the sequence for mTFP and actin 3' UTR. This construct also contained a Pac puromycin resistance gene for selection.

Transfection: Between 5-10 µg of each plasmid construct was transfected per reaction. Transfection and selection of plasmids in *M. brevicollis* was performed according to Woznica et al. (2021). Fluorescent cultures were clonally isolated, expanded, and frozen down.

### MbABCG1:SG imaging

To visualize the localization of MbABCG1::SG relative to the actin cortex and collar, we generated a MbABCG1::SG cell line co-expressing LifeAct::mCherry.

Late log-phase cells were aliquoted into 18-well chamber slide (Ibidi) for live imaging.

Cells were imaged on a Nikon AXR-NSPARC system equipped with a Plan Apochromat λ D 60x Water objective (NA – 1.2; WD - 0.18 mm). Fluorescent images were acquired using a FN25 galvano scanner, 488 nm and 568 nm lasers, and a 0.16 µM step size. At least 15 cells were imaged. Following acquisition, Z-stacks were 3D aligned and deconvolved using Nikon Elements software. Imaris software was used to visualize maximum intensity projections and 3D object rendering with transparency.

### MbABCG1:SG Flow Cytometry

MbABCG1::StayGold cells were split 1:20 the day before each experiment, washed, and resuspended in ASW at a density of 5-7 x $10^5$ cells/mL. Cells were aliquoted into 2-mL microcentrifuge tubes at a total volume of 500 µL per tube. Most treatments were carried out for 5 hours at RT and under gentle nutation. For the HNQ time course, compound was added to separate samples at staggered times so that all samples reached their respective endpoints simultaneously and could be acquired in a single run.

Deleted: prepared in technical replicates, and

Alkylquinolone treatments**:** Alkylquinolones (HHQ, 2-octyl-4-quinolone, HNQ, PQS, and 2-nonyl-3-hydroxy-4-quinolone) were added from DMSO stocks to indicated molar concentrations, standardized to 10 µL DMSO/tube.

CDN treatments: 2'3'-cGAMP and 3'3'-cGAMP were resuspended in water and applied to cells at a final concentration of 100 µM.

Conditioned media treatments: mPAO1 and *pqsA* conditioned media was prepared as described in above experiments. Cells were treated with 5% (vol/vol; 25 uL per 500 uL) conditioned media or an equivalent volume of LB as the control.

Deleted: strains were grown to an OD of 1.2, at which point cells were pelleted, and conditioned media was filter

**Acquisition and analysis.** Fifteen minutes before the 5-hour endpoint, samples were transferred as technical triplicates (120 uL each) into a tissue-culture-treated 96-well plate and acquired on a BD LSRFortessa flow cytometer. Detector voltages were set using untagged *M. brevicollis* such that choanoflagellate cells were resolved on the FSC/SSC plot (FSC, 350 V; SSC, 200 V; GFP, 442 V). A size gate was drawn around the bulk choanoflagellate population; no fluorescence gate was applied, as non-fluorescent (untagged) cells were run in parallel to define background. The median fluorescence intensity (MFI) of the StayGold/GFP channel (488 nm excitation) was quantified for the gated population.

**Graphing and Statistics**

Information about the quantification and statistical details of experiments can be found in the corresponding figure legends and methods. For most experiments, data were graphed and statistical analyses were performed using GraphPad Prism 11.0.0.

**Resource Availability**

Further information and requests for resources and reagents should be directed to Arielle Woznica -- woznica{at}utexas.edu.

**Material availability**

Plasmids and choanoflagellate cell lines used in this study are available upon request.

**Data and code availability**

RNA sequencing data files are available from the GEO repository #GSE355357.

Proteomics data: Data including FASTA files are available from the MassIVE data repository #MSV000103893

19. Powell, D. *Degust: Interactive RNA-Seq Analysis*.
